# ATR enforcement of the S/G2 checkpoint prevents premature S phase shutdown and genome instability

**DOI:** 10.64898/2026.05.07.723638

**Authors:** Melissa J. McEvoy, Joshua C. Saldivar

**Affiliations:** Cancer Early Detection Advanced Research Center, Knight Cancer Institute, Oregon Health & Science University, Portland, OR, USA; Program in Biomedical Sciences, School of Medicine, Oregon Health & Science University, Portland, OR, USA; Division of Oncological Sciences, Knight Cancer Institute, Oregon Health & Science University, Portland, OR, USA

## Abstract

The ATR-enforced S/G2 checkpoint activates during DNA replication to restrain CDK1-dependent phosphorylation of FOXM1 and subsequent transactivation of the G2/M gene network until the end of S phase. However, the extent to which this checkpoint ensures the completion of DNA replication and whether it safeguards genomic integrity has remained unknown. Here, we induce S/G2 checkpoint failure throughout S phase in non-malignant human epithelial cells using multiple ATR pathway inhibitors. Consequently, the mitotic kinase complex cyclin B1-CDK1 prematurely shuts-down the DNA replication program, preventing the completion of genome duplication. In turn, this leads to the retention of inactive replisomes on chromatin and unfired origins into the G2 phase, which induce subsequent accumulation of pan-nuclear ᵧH2AX and mitotic failure. Collectively, these findings indicate the S/G2 checkpoint ensures replication completion and genome stability.

## Introduction

Cell cycle checkpoints are essential regulatory mechanisms that ensure each phase of the cell cycle is complete before progression into the next phase, thereby mitigating DNA replication and chromosome segregation errors that could lead to genomic instability and cell death^1,2^. The mechanisms underlying the G1/S, G2/M, and metaphase-anaphase checkpoints are well established; however, the S/G2 checkpoint remains poorly understood and the central regulators of the S/G2 transition are under debate^3,4^. Moreover, the cellular consequences of checkpoint failure are unknown.

The S/G2 checkpoint is thought to monitor the completion of DNA replication and regulate entry into G2^3,4^. In its original description, the checkpoint was proposed to be enforced by the ATR-CHK1 (ataxia telangiectasia and Rad3-related; checkpoint kinase 1) pathway^3^. In the context of DNA damage, ATR is recruited to RPA-coated ssDNA, an intermediate that forms at stalled replication forks and resected DNA breaks and is subsequently activated by TOPBP1 and ETAA1^5–10^. Activated ATR triggers a phosphorylation cascade that results in the phosphorylation and activation of CHK1^11^. Once phosphorylated, CHK1 phosphorylates and inactivates the CDC25A-C (Cell Division Cycle 25 A-C) phosphatases, preventing the dephosphorylation of CDK1/2 at two inhibitory sites (T14 and Y15)^12^. Through this mechanism, ATR suppresses CDK1/2 activity and halts the cell cycle to allow DNA repair^13^.

While DNA damage amplifies ATR kinase activity, ATR is intrinsically active throughout each S phase, even in the absence of genotoxin-induced DNA damaged^3^. This intrinsic activity may be triggered by endogenous sources of DNA damage or a different ATR activating signal that is present during DNA replication. Regardless, ATR activity in S phase suppresses CDK1/2 activity^14^. Indeed, intrinsic ATR activity was shown to block CDK1-mediated phosphorylation and activation of the transcription factors FOXM1 (Forkhead Box Protein M1) and B-MYB throughout S phase^3,15,16^. In turn, this block prevented the expression of the G2/M gene network, including cyclin B1^3^ and consequently delayed the progression towards mitosis. Upon transition from S phase to G2 phase, ATR activity declines relieving the S/G2 checkpoint and allowing FOXM1-and B-MYB-dependent transactivation of the G2/M gene network and subsequent mitotic entry^3,15^.

Despite independent studies showing the ATR-CHK1 pathway delays FOXM1 activation, mitotic gene expression, and the onset of mitosis^3,4,15,17,18^, the extent to which ATR enforces the S/G2 checkpoint has remained disputed. For example, one study alternatively proposed that TRESLIN-MTBP monitors ongoing DNA replication and restrains progression into the G2 phase until replication is complete. Here, depletion of TRESLIN increased levels of cyclin B1 in S phase cells, suggesting that TRESLIN also regulates S/G2 transition^4^. Since TRESLIN-MTBP plays key roles in promoting origin firing^17,19,20^, it may serve to link origin firing to suppression of FOXM1, though the mechanistic details remain undefined.

A notable difference between TRESLIN loss and ATR inhibition is that when ATR is inhibited, there is a population of cells that begin to develop pan-nuclear ᵧH2AX, a DNA damage marker, in S phase^21^. While TRESLIN-deficient cells do exhibit a modest increase in ᵧH2AX, they lack the pan-nuclear staining seen with ATR inhibition^4^. Intriguingly, this striking staining pattern is not due to excessive origin firing, RPA exhaustion, or replication catastrophe^21^, phenotypes linked to ATR’s function in response chemically-induced replication stress^22^. Despite its prominence, the direct cause of pan-nuclear ᵧH2AX signaling and whether it is connected to failure of the S/G2 checkpoint remains unexplored.

Here, we reinforce the central role of ATR in regulating the S/G2 checkpoint and show that checkpoint inactivation triggers premature shutdown of the S phase replication program leaving replication origins unfired and inactive replisomes engaged with chromatin into the G2 phase. Consequently, cells fail to duplicate their DNA content, acquire pan-nuclear ᵧH2AX formation and undergo mitotic failure.

## Results

### ATR regulates the S/G2 checkpoint throughout S phase

ATR’s role as an enforcer of the S/G2 checkpoint was originally proposed following the observation that ATR inhibitors (ATRi) induced premature cyclin B1 expression in S phase^3^. However, this role of ATR has been questioned due to the relatively late accumulation of cyclin B1 near the end of S phase in ATR-inhibited cells^3^. Indeed, TRESLIN knockdown induces cyclin B1 expression throughout S phase, suggesting TRESLIN plays a greater S/G2 checkpoint role than ATR^4^. However, in these studies TRESLIN was knocked-down for 48 hours, whereas ATR was inhibited for only 6 hours. Thus, any conclusion on the importance of ATR vs TRESLIN in the S/G2 checkpoint is based on inequivalent experimental designs. Indeed, the difference in cyclin B1 accumulation in S phase may be driven by the duration of inactivation of the checkpoint rather than by the relative role of ATR vs TRESLIN in enforcing it.

For this reason, we extended the duration of ATR or CHK1 inhibition to 16 hours, and using quantitative image-based cytometry (QIBC)^22^, measured cyclin B1 levels across the cell cycle (Supplementary Fig 1a). The extended inhibition of ATR and CHK1 induced a prominent increase in cyclin B1 levels across S phase, and not only in late S phase, of non-malignant MCF10A cells (Fig. 1a and 1b and Supplementary Fig. 1b) and RPE-1 hTERT cells (Supplementary Fig. 1c), confirming the duration of ATR pathway inactivation determines the degree of cyclin B1 accumulation. Additionally, we noted that cells prematurely expressing cyclin B1 also showed decreases in EdU (5-ethynyl-2’-deoxyuridine) incorporation, indicating a reduction of replication and a potential premature exit from S phase (Fig. 1c - 1f and Supplementary Fig. 1d and 1e.

**Fig. 1:**
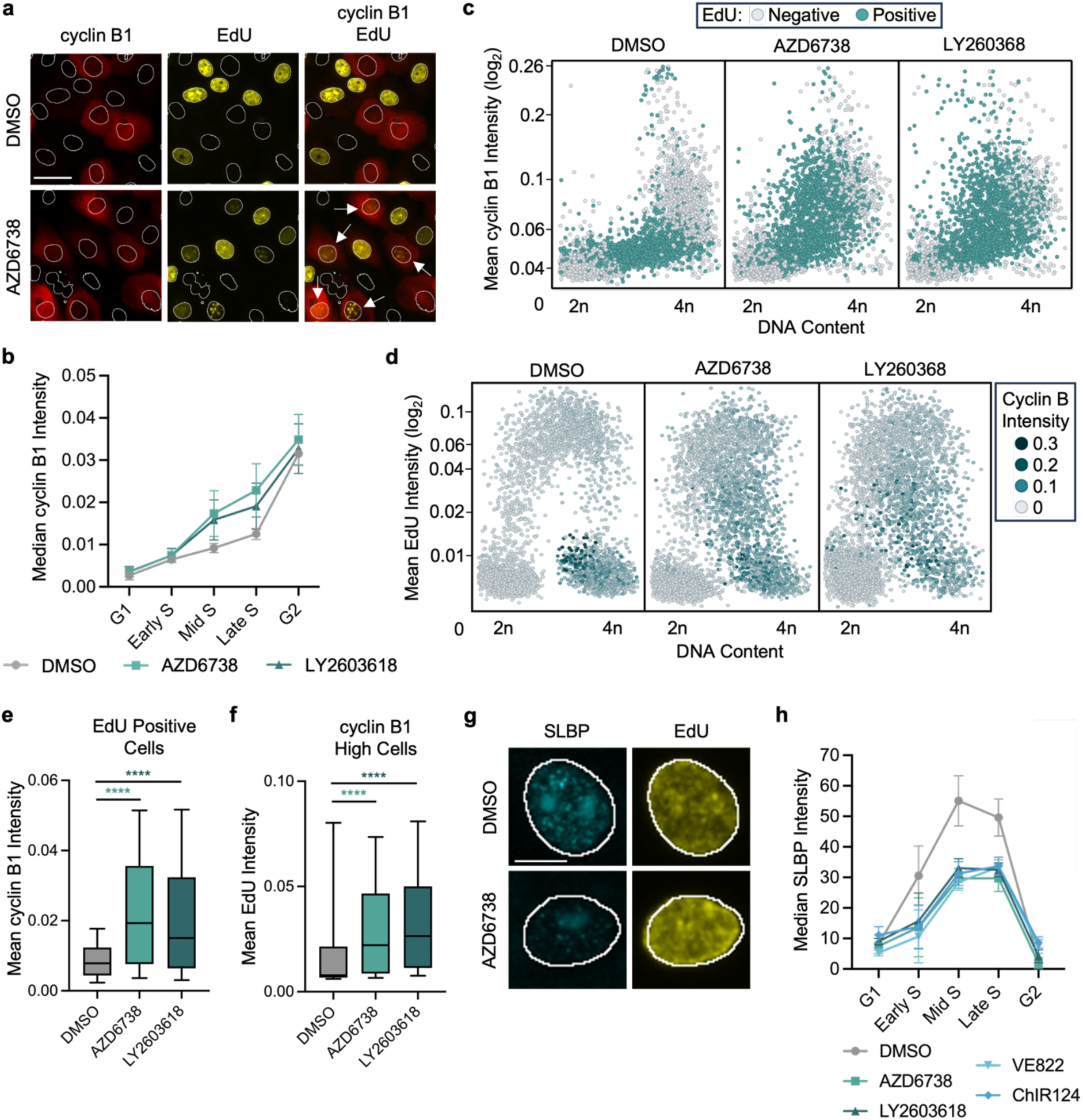
ATR regulates the S/G2 checkpoint throughout S phase. a.) Representative images of cyclin B1 and EdU in MCF10A cells treated with DMSO or 5μM AZD6738 for 16hrs. Representative of n=4 biological replicates. Arrows indicate nuclei positive for both EdU and cyclin B1. Scale bar is equal to 40μm. b.) Quantification of median cyclin B1 cytoplasmic intensity across the cell cycle in MCF10A cells treated with DMSO, 5μM AZD6738, or 2μM LY2603618 for 16hrs (n=4 biological replicates). Data are presented as mean ± SEM. c.) Scatterplots of DNA content (integrated DAPI intensity) versus mean cyclin B1 cytoplasmic intensity in MCF10A cells treated with DMSO, 5μM AZD6738, or 2μM LY2603618 for 16hrs. Each dot is representative of values from a singular cell. The plot is colored by EdU intensity (gating of negative/positive EdU values in Supplementary Fig. 1c) as denoted by the associated scale. Representative of n=4 biological replicates. d.) Scatterplots of DNA content (integrated DAPI intensity) versus mean EdU intensity in MCF10A cells treated with DMSO, 5μM AZD6738, or 2μM LY2603618 for 16hrs. Each dot is representative of values from a singular cell. The plot is colored by mean cyclin B1 intensity as denoted by the associated scale. Representative of n=4 biological replicates. e.) Quantification of mean cyclin B1 intensity values from Fig. 1c in EdU positive cells. Error bars are representative of 10-90 percentile range. A two-way analysis of variance (ANOVA) was performed to assess biological significance. *P* values: <0.0001 (DMSO versus AZD6738) and <0.0001 (DMSO versus LY2603618). f.) Quantification of mean EdU intensity values from Fig. 1d in cells with high cyclin B1 levels (gating of low/high cyclin B1 values in Supplementary Fig. 1b). Error bars are representative of 10-90 percentile range. *P* values: <0.0001 (DMSO versus AZD6738) and <0.0001 (DMSO versus LY2603618). g.) Representative images of SLBP and EdU in S phase MCF10A cells treated with DMSO or 5μM AZD6738 for 16hrs. Representative of n=3 biological replicates. Scale bar is equal to 10μm. h.) Quantification of median SLBP nuclear intensity across the cell cycle in MCF10A cells treated with DMSO, 5μM AZD6738, 5μM VE822, 2μM LY2603618, or 250nM ChIR124 for 1hr (n=3 biological replicates). Data are presented as mean ± SEM.

The need for an extended block of the checkpoint kinases is likely because cyclin B1 is a downstream marker of checkpoint failure, meaning multiple biochemical steps are required before accumulation can occur. The initial step of S/G2 checkpoint failure is CDK1 hyperactivation. Indeed, ATRi triggers CDK1-dependent phosphorylation of FOXM1 within 15 minutes throughout S phase^3,16^. As an independent marker of CDK1 hyperactivation and to evaluate ATR regulation of S phase processes, we measured Stem-Loop Binding Protein (SLBP), given that its levels peak in S phase and then decrease in G2 in a CDK1-dependent manner^23^. Importantly, within 1 hour of ATR and CHK1 inhibition, levels of SLBP dropped throughout S phase (Fig. 1g and 1h and Supplementary Fig. 1f and 1g). CDK1 inhibition, but not CDK2, rescued the ATRi-induced decrease in SLBP, restoring it to normal levels throughout S phase (Supplementary Fig. 1h and 1i). Together, these findings confirm that loss of ATR signaling perturbs the transition between S and G2 phase, both in terms of premature inactivation of S phase processes and premature expression of the G2/M gene network.

### Loss of the S/G2 checkpoint causes global DNA damage signaling

It is unknown whether ATR’s role in enforcing the checkpoint safeguards genome integrity and if it is essential for cell viability. We recently demonstrated that ATR inhibition triggers pan-nuclear γH2AX throughout S phase^21^. We hypothesize that the appearance of pan-nuclear γH2AX formation may be a consequence of premature cyclin B1 expression in S phase due to S/G2 checkpoint failure. Indeed, cyclin B1-CDK1 activity in S phase can induce DNA break formation due to premature disassembly of the replicative helicase^24^ and thus may underlie the formation of DNA damage upon ATR inhibition. To determine if this dramatic phenotype was a consequence of S/G2 checkpoint failure, we imaged γH2AX together with cyclin B1. Strikingly, the S phase cells with premature cyclin B1 expression were also positive for pan-nuclear γH2AX staining, indicating a strong correlation between S/G2 checkpoint failure and pan-nuclear DNA damage (Fig. 2a-2c 2d and Supplementary Fig. 2a). Furthermore, these pan-nuclear γH2AX-positive cells were the same population that exhibited reduced EdU incorporation, resembling cells exiting S phase (Fig. 1d). To determine if checkpoint failure induced DNA damage, we knocked-down cyclin B1 and measured γH2AX following ATR inhibition (Fig. 2d and Supplementary Fig. 1d and 2b). Importantly, cyclin B1 knockdown prevented the formation of pan-nuclear γH2AX staining throughout S phase, confirming S/G2 checkpoint failure is necessary for nucleus-wide DNA damage signaling.

**Fig. 2:**
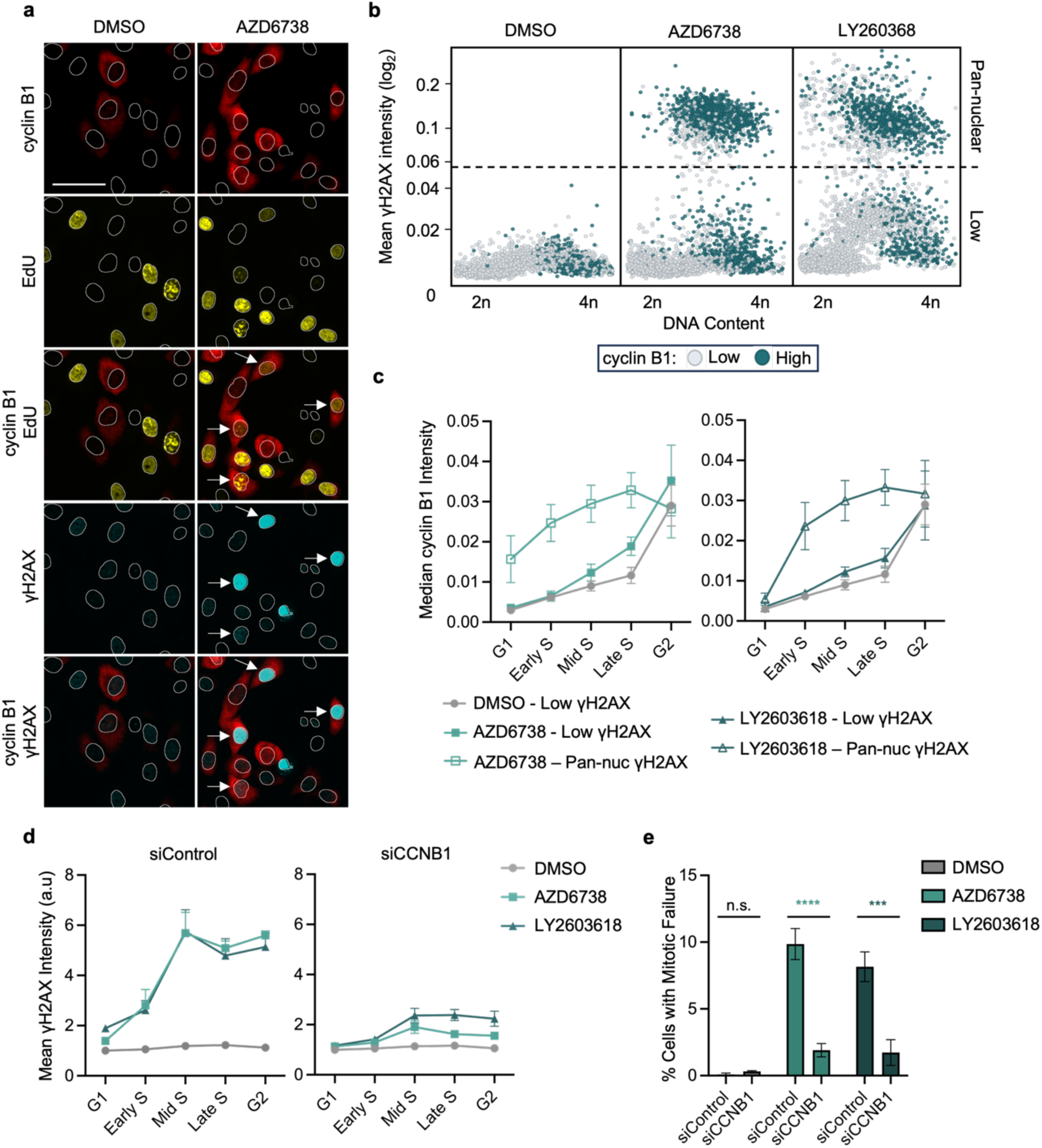
Loss of S/G2 checkpoint causes pan-nuclear DNA damage signaling. a.) Representative images of cyclin B1, EdU, and yH2AX in MCF10A cells treated with DMSO or 5μM AZD6738 for 16hrs. Representative of n=4 biological replicates. Arrows indicate nuclei positive for EdU, cyclin B1 and yH2AX. Scale bar is equal to 50μm. b.) Scatterplots of DNA content (integrated DAPI intensity) versus mean yH2AX intensity in MCF10A cells treated with DMSO, 5μM AZD6738, or 2μM LY2603618 for 16hrs. Each dot is representative of values from a singular cell. The plot is colored by high or low cyclin B1 intensity as denoted by the associated scale. Representative of n=4 biological replicates. c.) Quantification of median cyclin B1 cytoplasmic intensity in MCF10A cells with low or high yH2AX across the cell cycle. Cells were treated with DMSO, 5μM AZD6738, or 2μM LY2603618 for 16hrs (n=4 biological replicates). Data are presented as mean ± SEM. d.) Quantification of mean yH2AX nuclear intensities across the cell cycle from control and cyclin B1 knockdown MCF10A cells treated with DMSO, 5μM AZD6738, or 2μM LY2603618 for 16hrs (n=3 biological replicates). Data are presented as mean ± SEM. e.) Quantification of mitotic failure (defined in Supplementary Fig. 2e) in control and cyclin B1 knockdown MCF10A cells treated with DMSO, 5μM AZD6738, or 2μM LY2603618 for 16hrs (n=3 biological replicates). Percentages were calculated based on counts of 4 images per treatment for each biological replicate. A two-way analysis of variance (ANOVA) was performed to assess biological significance. *P* values: 0.9964 (siControl versus siCCNB1 + DMSO), <0.0001 (siControl versus siCCNB1 + AZD6738), and 0.0003 (siControl versus siCCNB1 + LY2603618). Data are presented as mean ± SEM.

To identify which DNA damage response kinase was mediating pan-nuclear DNA damage signaling in the absence of ATR activity, we inhibited ATM and DNA-PK while the ATR pathway was incapacitated. Notably, ATM inhibition did not change pan-nuclear γH2AX levels. By contrast, DNA-PK inhibition blocked the nucleus-wide spread of γH2AX (Supplementary Fig. 2c and 2d), suggesting ATRi-induced DNA damage leads to DNA-PK hyperactivation consistent with previous results showing DNA-PK acts as a back-up to loss of ATR^25,26^.

Additionally, we sought to identify whether S/G2 checkpoint failure compromised mitosis. Recently, we demonstrated that ATR inhibition induces mitotic slippage and mitotic catastrophe^21^, which we here refer to under the umbrella term of mitotic failure (Supplementary Fig. 2e). To determine if premature expression of cyclin B1 drives mitotic failure, we knocked down cyclin B1 and quantified the percentage of cells that underwent mitotic failure when the ATR-CHK1 pathway was perturbed (Fig. 2e). Cyclin B1 knockdown reduced the population of cells with mitotic failure, indicating that the premature expression of cyclin B1 induced by ATR and CHK1 inhibition not only compromises genomic stability, but also cellular viability.

### S/G2 checkpoint failure prevents completion of DNA replication

Next, as ATR and CHK1 inhibition decreases EdU incorporation in S phase cells with elevated cyclin B1 levels (Fig. 1d), we reasoned S/G2 checkpoint failure may block replication progression leading to incomplete replication upon transition to the G2 phase. To this end, we focused our analysis on EdU-negative cells with a DNA content of ≥3n, a population that by QIBC analysis corresponds to the G2 phase (Supplementary Fig. 3a). In this population of cells, we analyzed the levels of replisome components on chromatin following pre-extraction prior to fixation and QIBC. As expected, in control cells, the PCNA (proliferating cell nuclear antigen) clamp was only present on chromatin in S phase cells actively replicating DNA (EdU positive cells) and not in the G2 population (Fig. 3a and 3b). Notably, ATR- and CHK1-inhibited cells with pan-nuclear γH2AX exhibited a marked increase in PCNA in the EdU-negative population, indicating replisomes are engaged with chromatin yet inactive (Fig. 3a and 3b and Supplementary Fig. 3b and 3c), and these inactive replisomes were retained on chromatin into the G2 phase (Fig. 3b and Supplementary Fig. 3c. Similarly, the replicative helicase subunit MCM2 was also observed to be retained on chromatin into the G2 phase following ATR or CHK1 inhibition (Fig. 3c and 3d and Supplementary Fig. 3d and 3e).

**Fig. 3:**
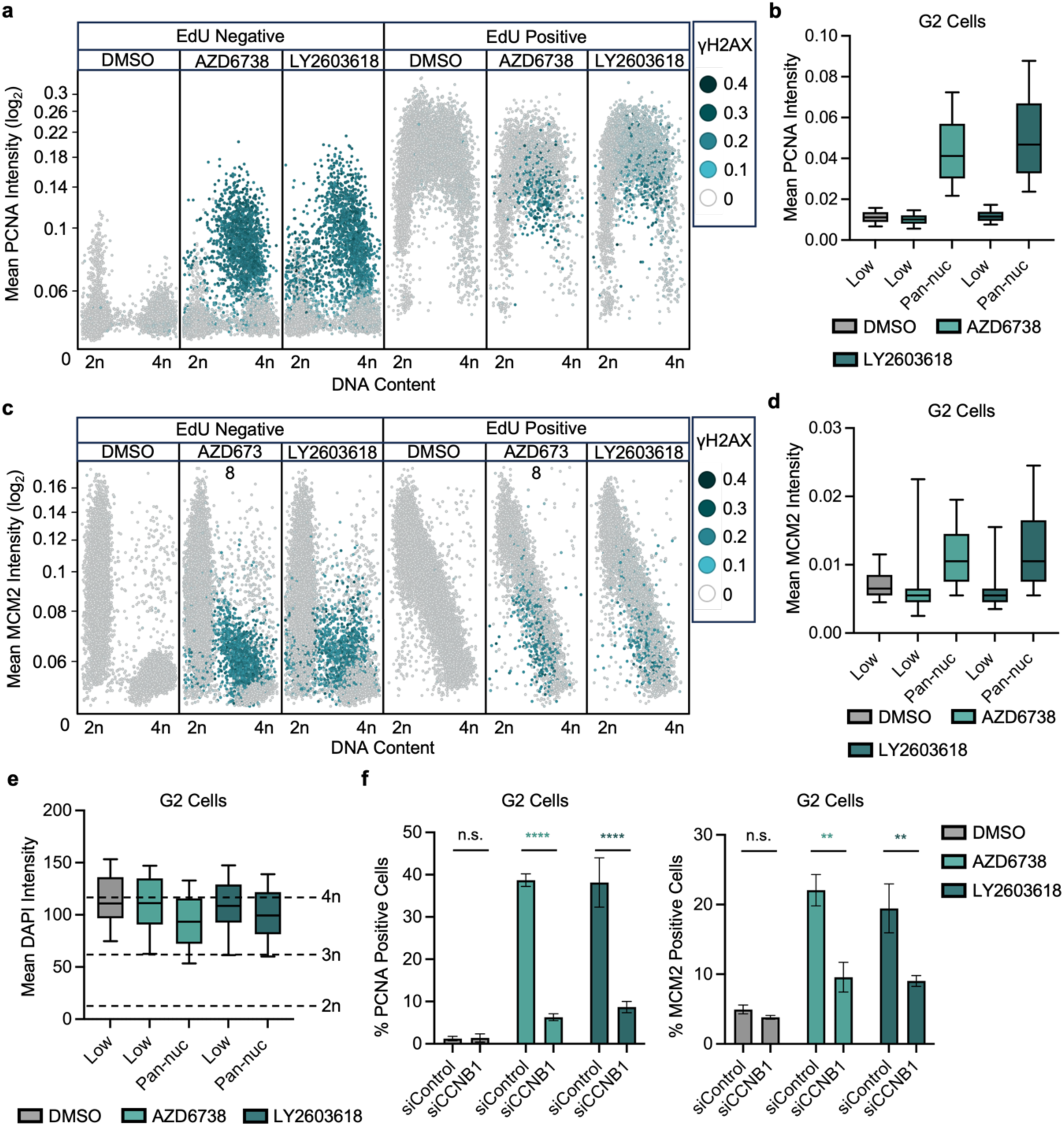
S/G2 checkpoint failure prevents completion of DNA replication. a.) Scatterplots of mean chromatin-bound PCNA intensity versus DNA content in pre-extracted MCF10A cells treated with DMSO, 5μM AZD6738, or 2μM LY2603618 for 16hrs. Plots are divided by EdU negative/positive cells and by treatment. The plot is colored by mean yH2AX intensity as denoted by the associated scale. Representative of n=3 biological replicates. b.) Quantification of mean PCNA intensities from Fig. 3a in pre-extracted MCF10A cells with low or high yH2AX in the defined G2 population (Supplemental Fig. 3a). Error bars are representative of 10-90 percentile range. c.) Scatterplots of mean MCM2 intensity versus DNA content in pre-extracted MCF10A cells treated with DMSO, 5μM AZD6738, or 2μM LY2603618 for 16hrs. Plots are divided by EdU negative/positive cells and by treatment. The plot is colored by mean yH2AX intensity as denoted by the associated scale. Representative of n=3 biological replicates. d.) Quantification of mean MCM2 intensities from Fig. 3c in pre-extracted MCF10A cells with low or high yH2AX in the defined G2 population. Error bars are representative of 10-90 percentile range. e.) Quantification of integrated DAPI intensities to measure DNA content in pre-extracted MCF10A cells with low or high yH2AX in the defined G2 population (n=3 biological replicates). Error bars are representative of 10-90 percentile range. f.) Quantification of the percentage of PCNA (left) or MCM2 (right) positive pre-extracted MCF10A cells (gating of PCNA and MCM2 shown in Supplementary Fig. 3b and 3c) in the defined G2 population of control and cyclin B1 knockdown MCF10A cells treated with DMSO, 5μM AZD6738, or 2μM LY2603618 for 16hrs (n=3 biological replicates). A two-way analysis of variance (ANOVA) was performed for each plot to assess biological significance. *P* values for PCNA plot: >0.9999 (DMSO siControl versus siCCNB1), <0.0001 (AZD6738 siControl versus siCCNB1), and <0.0001 (LY2603618 siControl versus siCCNB1). *P* values for MCM2 plot: 0.9694 (DMSO siControl versus siCCNB1), 0.0021 (AZD6738 siControl versus siCCNB1), and 0.0081 (LY2603618 siControl versus siCCNB1). Data are presented as mean ± SEM.

Next, we quantified the DNA content of the G2 population with S/G2 checkpoint failure and pan-nuclear γH2AX. This analysis revealed a failure to fully duplicate the DNA content following ATR or CHK1 inhibition, with the most pronounced decrease observed in cells exhibiting pan-nuclear γH2AX staining (Fig. 3e). These data suggest that S/G2 checkpoint failure allows transition into the G2 phase despite DNA replication being incomplete, resulting in the retention of inactive replisome components on chromatin into the G2 phase.

Finally, to determine if the retention of replisome components on chromatin into the G2 phase was a result of the early expression of G2/M gene network, we knocked-down cyclin B1 and measured the percentage of G2 cells positive for PCNA or MCM2 (Supplementary Fig. 3b and 3d). Importantly, cyclin B1 knockdown significantly reduced the percentage of cells with PCNA and MCM2 retained on chromatin in G2 (Fig. 3f). These results demonstrate that untimely cyclin B1 expression in S phase causes the premature loss of replication and the inappropriate persistence of replisome factors on chromatin in G2.

### S/G2 checkpoint failure causes progressive shutdown of the replication program throughout S phase

Upon observing the persistence of PCNA and MCM2 on chromatin in the G2 phase, we asked if this was due to replisome inactivation, a failure to fire replication origins, or both. To address this, we first sought to define the point in the replication program when S/G2 checkpoint failure leads to loss of DNA replication. Confocal fluorescence imaging of MCM2 and PCNA in ATR- or CHK1-inhibited cells revealed distinct staining patterns indicative of particular stages in the replication program. Specifically, in the population of EdU positive cells with pan-nuclear DNA damage, PCNA chromatin binding had very pronounced early-to-mid S phase staining patterns (Fig. 4a and Supplementary Fig. 4a). Low levels of EdU incorporation in these cells suggest most replisomes were either functioning very inefficiently or were inactivated. Cells with pan-nuclear DNA damage but no detectable EdU incorporation retained PCNA in a late S phase pattern. Here, the presence of PCNA and the lack of EdU incorporation suggest a complete shutdown of the replication program despite incomplete replication. Similar to PCNA, MCM2 staining in the population of cells with pan-nuclear γH2AX, revealed early-to-mid S phase patterns in cells with low EdU incorporation and a late S pattern in the EdU-negative cells (Fig. 4b and Supplementary Fig. 4b). Thus, we propose that loss of the ATR-enforced checkpoint triggers progressive loss of replication throughout S phase and a complete shutdown of the replication program by late S phase.

**Fig. 4:**
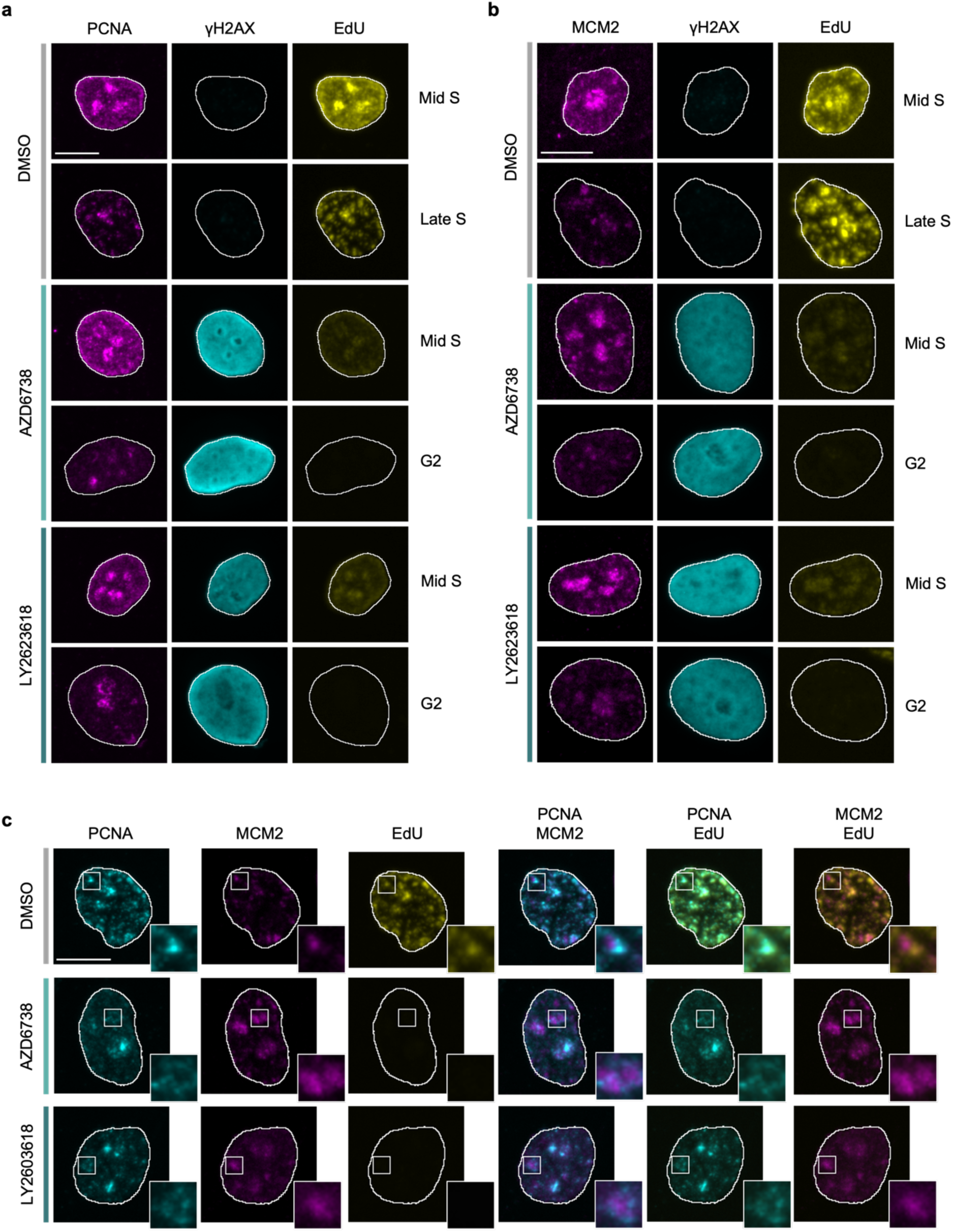
Incomplete replication results from progressive shut down of the replication program throughout S phase. a.) Representative images of PCNA, yH2AX, and EdU in pre-extracted MCF10A nuclei treated with DMSO, 5μM AZD6738, or 2μM LY2603618 for 16hrs. Cell cycle status is indicated in the figure. Representative of n=3 biological replicates. Scale bar is equal to 10μm. b.) Representative images of MCM2, yH2AX, and EdU in pre-extracted MCF10A nuclei treated with DMSO, 5μM AZD6738, or 2μM LY2603618 for 16hrs. Cell cycle status is indicated in the figure. Representative of n=3 biological replicates. Scale bar is equal to 10μm. C.) Representative images of PCNA, MCM2, and EdU in pre-extracted MCF10A nuclei treated with DMSO, 5μM AZD6738, or 2μM LY2603618 for 16hrs. The DMSO treated nuclei shows a cell in late S phase, while the nuclei from the other treatments show EdU negative cells that have halted replication. Whit boxes indicate areas of the nuclei that have been magnified. Representative of n=3 biological replicates. Scale bar is equal to 10μm.

Next, we co-stained PCNA and MCM2 to see if they colocalize when retained on chromatin in the G2 phase. When colocalized with PCNA, MCM2 is thought to mark the replicative MCM helicase as it is in close proximity to PCNA and the rest of the replisome^27^. MCM2 signals that do not colocalize with PCNA indicate the presence of licensed but unfired origins, consistent with the established model in which MCM2 is evicted from chromatin in a replication-dependent manner. Of note, we observed that in the G2 cells with retained PCNA and MCM2, there was limited co-localization between the two markers, with the majority of MCM2 staining appearing independently of PCNA (Fig. 4c). PCNA staining was often near MCM2 but organized in focal regions neighboring MCM2-enriched regions. The presence of PCNA foci in EdU-negative cells suggest replisomes are no longer functional despite being retained. The presence of MCM2 on neighboring chromatin indicates these G2 cells have unfired origins, a hallmark of under-replication and a sign of premature replication shutdown^28^. Together, these results suggest that throughout S phase, loss of the S/G2 checkpoint triggers a progressive shutdown of the replication program, inactivating mid-to-late S phase replisomes and leaving late origins unfired. Consequently, cells fail to fully duplicate their DNA content before exiting S phase.

### Pan-nuclear DNA damage is caused by persistent replisome engagement with chromatin in the G2 phase

Given our data that premature cyclin B1 expression is necessary for both premature shutdown of the replication program and pan-nuclear γH2AX, we reasoned that these two detrimental phenotypes were causally linked. Cyclin B1-CDK1 expression in S phase leads to DNA break formation due to TRAIP-dependent ubiquitination and removal of the replicative helicase^24^. The process of TRAIP-dependent ubiquitination is counterbalanced by the deubiquitinase USP37, which stabilizes replisome components and prevents their unscheduled disassembly, particularly under conditions of replication stress^29,30^. Using this replisome removal pathway as a tool to study replisome retention in ATR- and CHK1-inhibited cells, we targeted TRAIP and USP37 individually with siRNAs and measured chromatin-bound PCNA and MCM2. We predicted that knockdown of USP37 would allow TRAIP-dependent clearance of replisomes from chromatin in cells with S/G2 checkpoint failure. Consistent with our prediction, USP37, but not TRAIP, knockdown strongly decreased the retention of PCNA and MCM2 on chromatin in G2 cells (Fig. 5a-5d).

**Fig. 5:**
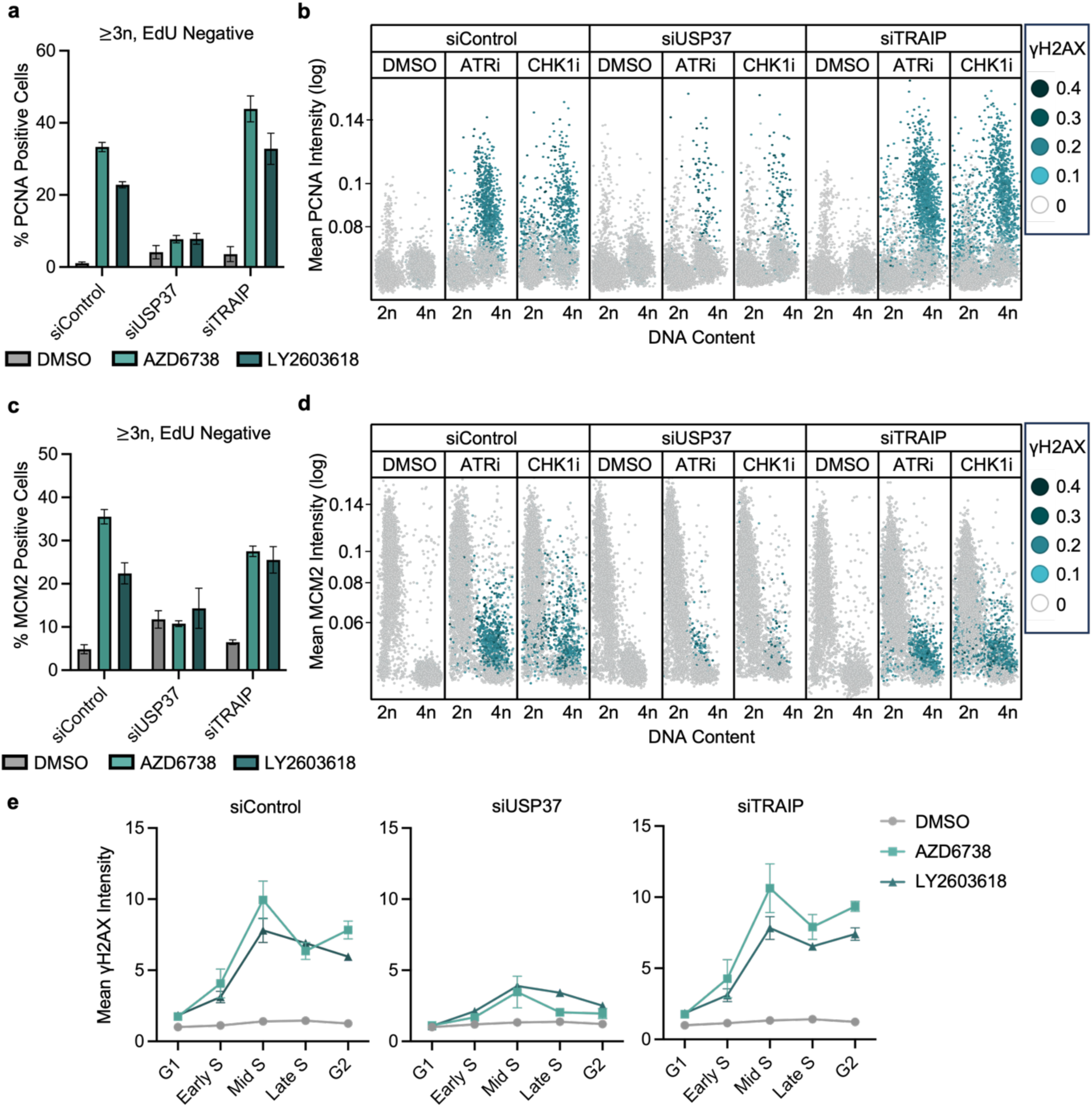
Pan-nuclear DNA damage is caused by persistent replisome engagement with chromatin in the G2 phase. a.) Quantification the percentage of PCNA positive pre-extracted cells in the defined G2 population of control, USP37, or TRAIP knockdown MCF10A cells treated with DMSO, 5μM AZD6738, or 2μM LY2603618 for 16hrs (n=3 biological replicates). A two-way analysis of variance (ANOVA) was performed to assess biological significance. *P* values: 0.5397 (DMSO siControl versus siUSP37), 0.6511 (DMSO siControl versus siTRAIP), <0.0001 (AZD6738 siControl versus siUSP37), 0.0068 (AZD6738 siControl versus siTRAIP), 0.0003 (LY2603618 siControl versus siUSP37), and 0.0101 (LY2603618 siControl versus siUSP37). Data are presented as mean ± SEM. b.) Scatterplots of mean PCNA intensity versus DNA content in control, USP37, or TRAIP knockdown and pre-extracted MCF10A cells treated with DMSO, 5μM AZD6738, or 2μM LY2603618 for 16hrs. Plots are divided by EdU negative/positive cells and by treatment. The plot is colored by mean yH2AX intensity as denoted by the associated scale. Representative of n=3 biological replicates. c.) Quantification of the percentage of MCM2 positive pre-extracted cells in the defined G2 population of control, USP37, or TRAIP knockdown MCF10A cells treated with DMSO, 5μM AZD6738, or 2μM LY2603618 for 16hrs (n=3 biological replicates). A two-way analysis of variance (ANOVA) was performed to assess biological significance. *P* values: 0.0837 (DMSO siControl versus siUSP37), 0.8341 (DMSO siControl versus siTRAIP), <0.0001 (AZD6738 siControl versus siUSP37), 0.0433 (AZD6738 siControl versus siTRAIP), 0.0410 (LY2603618 siControl versus siUSP37), and 0.5357 (LY2603618 siControl versus siUSP37). Data are presented as mean ± SEM. d.) Scatterplots of mean MCM2 intensity versus DNA content in control, USP37, or TRAIP knockdown and pre-extracted MCF10A cells treated with DMSO, 5μM AZD6738, or 2μM LY2603618 for 16hrs. Plots are divided by EdU negative/positive cells and by treatment. The plot is colored by mean yH2AX intensity as denoted by the associated scale. Representative of n=3 biological replicates. e.) Quantification of mean yH2AX nuclear intensities across the cell cycle from control, USP37, or TRAIP knockdown and pre-extracted MCF10A cells treated with DMSO, 5μM AZD6738, or 2μM LY2603618 for 16hrs (n=3 biological replicates). Data are presented as mean ± SEM.

Next, we asked whether the retention of inactive replisome components on chromatin in the G2 phase was necessary for the nucleus-wide DNA damage. We hypothesized this could be possible as a previous study in yeast showed the retention of PCNA in G2 induced by the deletion of *elg1,* a gene that encodes for a PCNA unloading factor, to be linked to genome instability^31^. Thus, we quantified γH2AX following USP37 or TRAIP depletion, and importantly, USP37 knockdown drastically reduced the formation of pan-nuclear γH2AX. In contrast, TRAIP knockdown slightly heightened pan-nuclear γH2AX accumulation (Fig. 5e). Collectively, these results demonstrate that the retention of replisome components on chromatin after S phase exit contributes to pan-nuclear DNA damage in S/G2 checkpoint defective cells.

### ATR inhibitor-resistant cells maintain genome stability under ATRi

ATR is essential for viability of proliferating cells, yet whether ATR’s essential function is linked to the S/G2 checkpoint is unknown. To address this question, we developed an ATR inhibitor-resistant MCF10A cell line via prolonged culturing of parental cells with gradually increasing concentrations of ATRi (Fig. 6a). This cell line, henceforth referred to as ATRi-R, can proliferate during prolonged culturing in the presence of full ATR inhibition, unlike naïve MCF10As, which fail to proliferate and gradually die off when cultured in the same conditions (Fig. 6b). Consistent with this resistance, ATRi-R cells are largely impervious to the induction of pan-nuclear γH2AX following ATR or CHK1 inhibition (Fig. 6c and 6d), indicating these cells have either restored the pathway downstream of CHK1 or rely on a secondary back-up pathway.

**Fig. 6:**
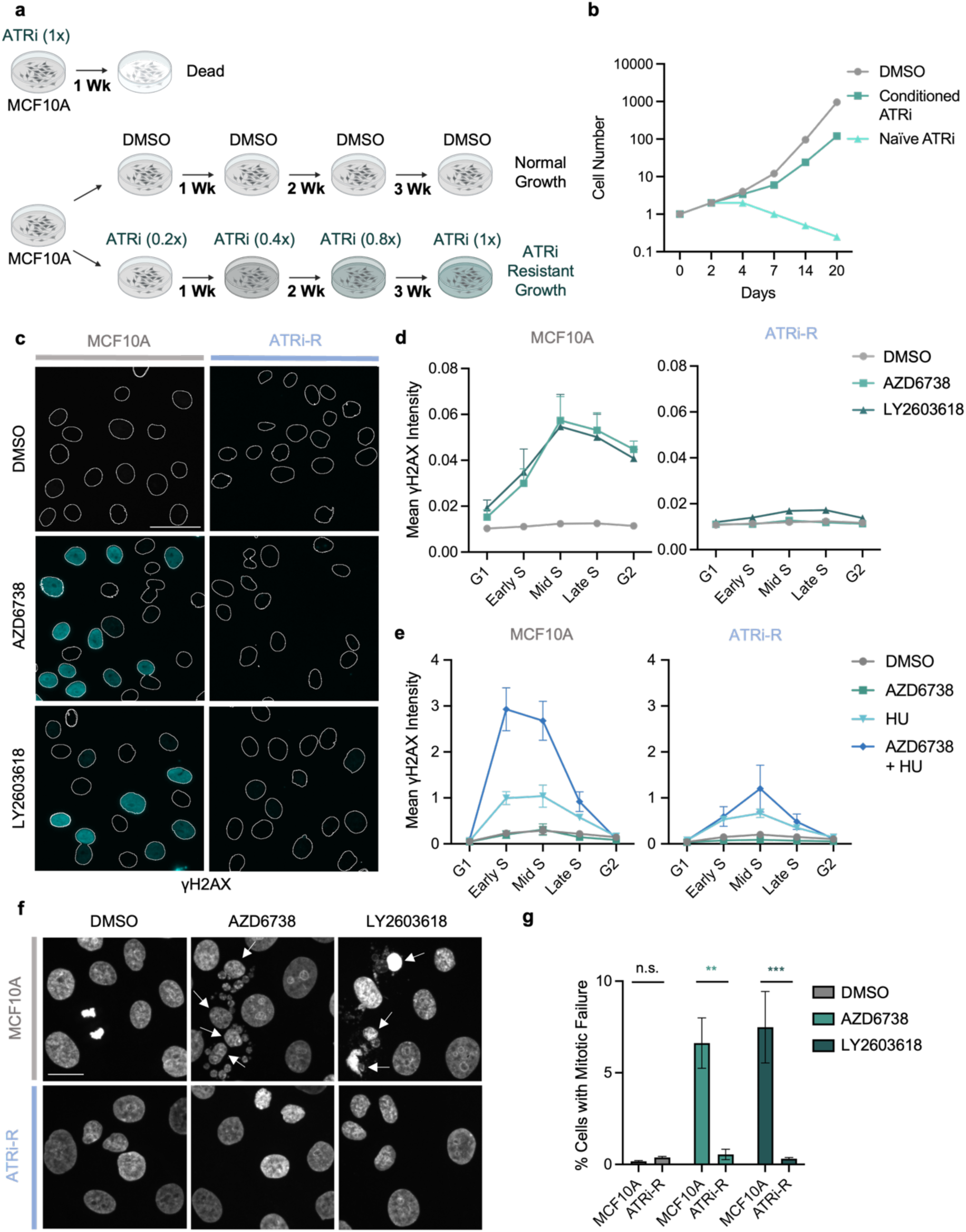
ATR inhibitor-resistant cells maintain genome stability under ATRi. a.) Schematic of ATRi resistant cell line development (ATRi-R). b.) Cell proliferation plot of ATRi-R cells compared to controls under 5μM AZD6738 treatment. c.) Representative images of yH2AX in MCF10A and ATRi-R cells treated with DMSO, 5μM AZD6738, or 2μM LY2603618 for 16hrs. Representative of n=3 biological replicates. Scale bar is equal to 50μm. d.) Quantification of mean yH2AX nuclear intensity across the cell cycle from MCF10A and ATRi-R cells treated with DMSO, 5μM AZD6738, or 2μM LY2603618 for 16hrs (n=3 biological replicates). Data are presented as mean ± SEM. e.) Quantification of mean yH2AX nuclear intensity across the cell cycle from MCF10A cells treated with DMSO, 5μM AZD6738, or 1mM hydroxyurea or a combination of 5μM AZD6738 and 1mM hydroxyurea 1.5hrs (n=3 biological replicates). Data are presented as mean ± SEM. f.) Representative images of DAPI stained nuclei in MCF10A and ATRi-R cells treated with DMSO, 5μM AZD6738, or 2μM LY2603618 for 16hrs. Representative of n=3 biological replicates. Arrows indicate cells with mitotic slippage. Scale bar is equal to 20μm. g.) Quantification of mitotic failure (defined in Supplementary Fig. 2C) in control and cyclin B1 knockdown MCF10A cells treated with DMSO, 5μM AZD6738, or 2μM LY2603618 for 16hrs (n=3 biological replicates). A two-way analysis of variance (ANOVA) was performed to assess biological significance. *P* values: 0.9983 (DMSO MCF10A versus ATRi-R), 0.0027 (AZD6738 MCF10A versus ATRi-R), 0.0007 (LY2603618 MCF10A versus ATRi-R). Data are presented as mean ± SEM.

We tested if either ATM or DNA-PK was protective to ATRi-R S phase cells. ATM inhibition had no impact on DNA damage levels across S phase (Supplementary Fig. 5a). By contrast, DNA-PK inhibition induced a moderate increase in DNA damage in early S phase cells, irrespective of whether ATR or CHK1 was also inhibited (Supplementary Fig. 5b). This observation is consistent with previous observations that DNA-PK substitutes for loss of ATR activity.

We next sought to determine if ATRi-R cells were resistant to replication protein A (RPA) exhaustion induced by combined ATR inhibition and hydroxyurea-induced fork stalling. Using γH2AX formation as a readout for exhaustion, ATRi-R cells displayed only a modest reduction in γH2AX signaling relative to MCF10A cells in response to hydroxyurea alone. In contrast, ATRi-R cells exhibited a pronounced rescue when treated with hydroxyurea in combination with ATR inhibition, a condition that robustly induces RPA exhaustion in parental cells (Fig. 6e). These results indicate that ATRi resistance selectively mitigates replication stress phenotypes that are exacerbated by ATR pathway inhibition. Finally, we found that ATR-R cells could rescue ATRi and CHK1i induced mitotic failure seen in parental MCF10A cells (Fig. 6f-6g).

### ATRi-R cells restore essential functions of the ATR-enforced S/G2 checkpoint

Upon confirming ATRi-R cells rescued cellular viability in ATR-inhibited cells, we sought to uncover whether the S/G2 checkpoint was restored in these cells. Quantification of cyclin B1 levels throughout the cell cycle in ATRi-R cells treated with ATRi and CHK1i revealed that inhibition of these kinases did not induce the premature expression of cyclin B1 in S phase cells (Fig. 7a and 7b). Next, we quantified SLBP levels across the cell cycle and observed increased SLBP expression in ATRi-R cells (Fig. 7c). ATR and CHK1 inhibition still induced a reduction in SLBP levels during S phase in ATRi-R cells; however, the magnitude of this decrease was attenuated and SLBP were comparable to those observed in untreated parental MCF10A cells. Finally, analysis of pFOXM1 levels in the ATRi-R cells revealed they were largely resistant to ATRi-induced phosphorylation of FOXM1 in S phase (Fig. 7d and 7e). Together, these data indicate that the S/G2 checkpoint was restored in the ATRi-resistant cell line, suggesting ATR enforcement of S/G2 checkpoint is essential for maintaining cellular viability.

**Fig. 7:**
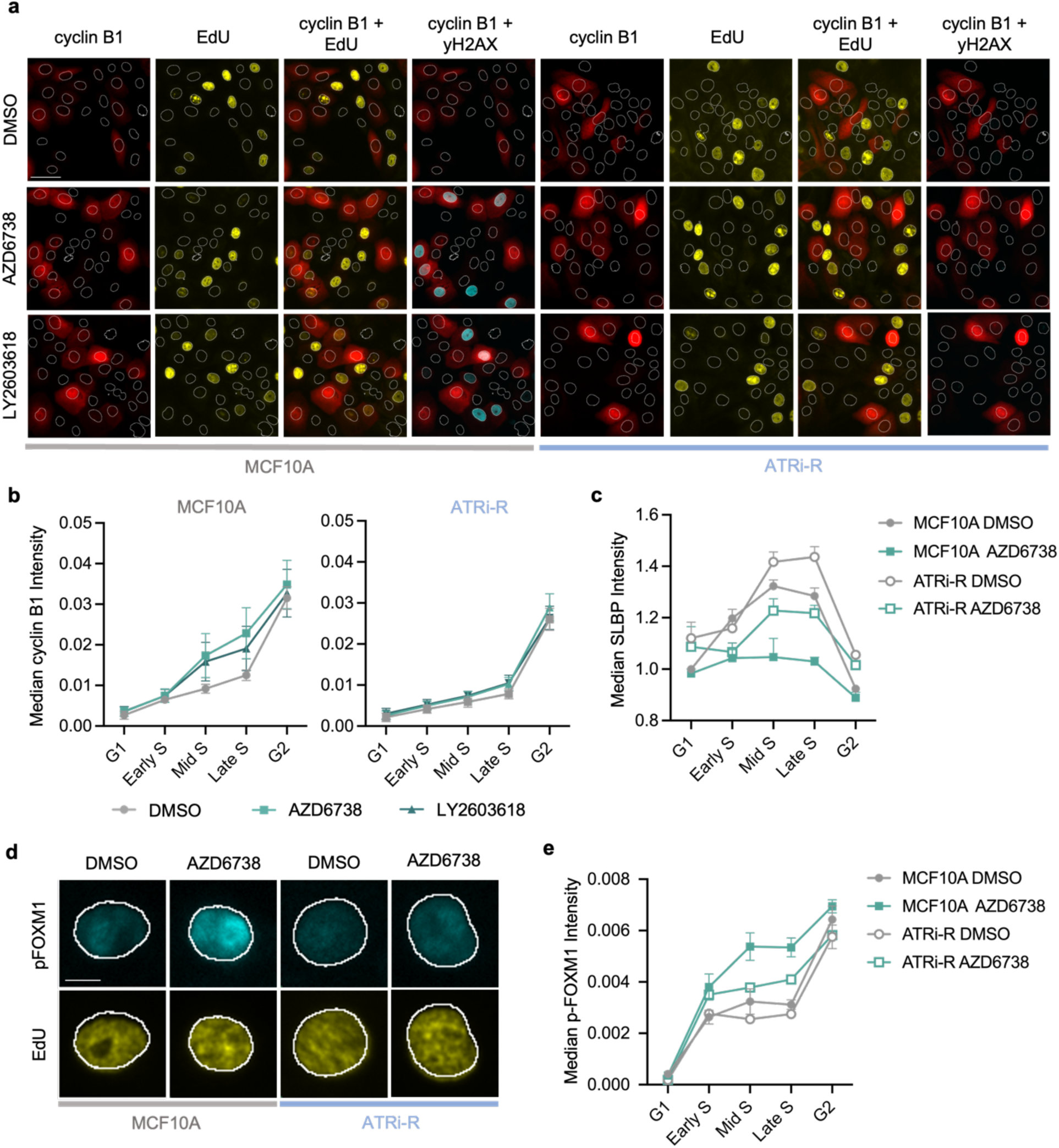
Essential S/G2 checkpoint functions are restored in ATRi resistant cells. a.) Representative images of cyclin B1, EdU, and yH2AX in MCF10A and ATRi-R cells treated with DMSO or 5μM AZD6738 for 16hrs. Representative of n=4 biological replicates. Scale bar is equal to 50μm. b.) Quantification of median cyclin B1 cytoplasmic intensity across the cell cycle from MCF10A cells treated with DMSO, 5μM AZD6738, or 2μM LY2603618 for 16hrs (n=4 biological replicates). Note: MCF10A data was also shown in Fig.1b. Data are presented as mean ± SEM. c.) Representative images of pFOXM1 (T600) and EdU in MCF10A and ATRi-R cells treated with DMSO or 5μM AZD6738 for 1hr. Representative of n=3 biological replicates. Scale bar is equal to 10μm. d.) Quantification of median pFOXM1 (T600) nuclear intensity across the cell cycle from MCF10A cells treated with DMSO, 5μM AZD6738, or 2μM LY2603618 for 1hr (n=3 biological replicates). Data are presented as mean ± SEM. e.) Quantification of median SLBP nuclear intensity across the cell cycle from MCF10A and ATRi-R cells treated with DMSO, 5μM AZD6738, or 2μM LY2603618 for 1hr (n=3 biological replicates). Data are presented as mean ± SEM.

## Discussion

We propose that enforcement of the S/G2 checkpoint is an essential function of ATR required to couple replication completion with orderly progression into G2. In contrast to a recent model stating ATR acts primarily as a brake to slow progression through S phase, we find that ATR activity is continuously required throughout S phase to prevent the S/G2 transition. Inhibition of ATR-CHK1 signaling results in a complete collapse of S/G2 checkpoint control, evidenced by premature SLBP degradation, early cyclin B1 accumulation, and replication shutdown during S phase. Thus, rather than simply monitoring the pace of cell-cycle progression, ATR actively suppresses untimely CDK1 activation until genome duplication is complete.

The consequences of ATR-CHK1 related S/G2 checkpoint failure extend beyond the early initiation of the G2 transcriptional program. We observe that checkpoint failure uncouples initiation of G2 processes from replication completion, leading to premature shutdown of the DNA replication program and entry into G2 with under-replicated genomes. This premature shutdown may explain why ATR inhibition induces common fragile site expression^32–34^, a broken chromosome phenotype in mitotic cells caused by incomplete DNA replication^35,36^.

As a consequence of replication shutdown, cells enter G2 with unfired replication origins and inactive replisomes that persist on chromatin despite having exited S phase. Notably, the spatial patterns of EdU and chromatin-bound PCNA and MCM2 suggest the replication program undergoes a gradual shutdown beginning in early S phase and becoming fully inactive in late S phase. These findings suggest that ATR, via S/G2 checkpoint enforcement, ensures the completion of DNA replication prior to the initiation of G2. Given that these detrimental phenotypes are dependent on cyclin B1 expression in S phase, we propose the mitotic CDK activity that normal builds up in G2 is incompatible with the replication program. In this way, cells may have an additional mechanism to prevent endoreplication in the G2 phase prior to mitotic division.

Another striking consequence of checkpoint failure upon inhibition of the ATR-CHK1 pathway is the formation of pan-nuclear γH2AX. This global DNA damage phenotype is triggered by premature cyclin B1 expression in S phase. Previous studies have shown the cyclin B1-CDK1 activity in S phase triggers replisome disassembly in TRAIP dependent manner^24^. Moreover, forced clearance of chromatin-engaged replisomes via USP37 knockdown rescues pan-nuclear DNA damage accumulation, linking the premature shutdown of S phase to nucleus-wide DNA damage. These observations argue that the widespread γH2AX observed following ATR inhibition is not simply a direct consequence of fork stalling but rather reflects a catastrophic loss of coordination between replication completion and cell-cycle progression. Moreover, this loss of coordination results in severe mitotic defects, including mitotic slippage. Intriguingly, we recently showed both pan-nuclear γH2AX and mitotic slippage were dependent on Mediator complex subunit 1 (MED1)^21^. Both MED1 and ATR colocalize within the histone locus body (HLB), and it is within this compartment that ATR suppresses CDK1. We speculate the HLB may be a hub of S/G2 checkpoint enforcement, though ATR may also suppress CDK1 throughout the nucleus.

In addition to redefining the conceptual framework of the S/G2 checkpoint, our findings also provide important implications for clinical usage of ATR inhibitors. ATR inhibitors are currently in late-stage clinical trials as cancer therapeutics, largely based on the premise that cancer cells are particularly dependent on ATR to tolerate elevated replication stress^37,38^. Our results suggest that ATR inhibition exerts its cytotoxic effects not only by exacerbating replication stress, but also by dismantling the S/G2 checkpoint, thereby forcing cells into a state in which CDK1 activation, replication progression, and mitotic gene expression become uncoupled. This unlocks the potential for novel treatment plans that may increase the efficacy of ATR inhibitors. Combination strategies that further destabilize checkpoint enforcement or promote premature CDK activation may potentiate the effects of ATR inhibitors, either alone or alongside existing regimens. It is also evident from our results that S/G2 checkpoint restoration may be a potential mechanism for developed resistance to ATR inhibitors, as our ATR inhibitor resistant cells restored S/G2 checkpoint functionality. Future research uncovering the mechanism through which cells restore the S/G2 checkpoint in ATR-inhibited cells may lead to novel combination therapies to improve the efficacy of ATR inhibitors.

Finally, our data imply that the replication program and activation of mitotic kinases are mutually exclusive: DNA replication blocks cyclin B1 expression and cyclin B1 expression blocks the replication program. Through this mutual exclusivity, the temporal order of the cell cycle is maintained ensuring replication is complete prior to mitosis, without which, cell viability would be lost. This elegant means to control the temporal order is possible, in part, through the workings of the ATR-enforced S/G2 checkpoint.

## Methods

### Cell Culture

Human MCF10A cells (ATCC, CRL-10317) were grown in DMEM/F12 medium (ThermoFisher Scientific, 11330057) supplemented with 100U/mL penicillin/streptomycin (Thermo Fisher Scientific, 15140122), 5% horse serum (ThermoFisher Scientific, 16050122), 20ng/ml epidermal growth factor (EGF) (Sigma-Aldrich, E5036), 0.5 mg/mL hydrocortisone, 10μg/mL insulin (Sigma-Aldrich, I1882-100MG), and 100ng/mL cholera toxin (Sigma-Aldrich, C8052). Human hTERT RPE-1 cells (ATCC, CRL-4000) were grown in DMEM/F12 medium supplemented with 10% fetal bovine serum (Thermo Fisher Scientific, A5256801), 100U/mL penicillin/streptomycin, and 10µg/mL hygromycin B (Roche, 10843555001). All cells were grown in a humidified atmosphere at 37°C with 5% CO_2_.

### ATRi-R Cell Line Development

The ATR inhibitor resistant MCF10A cell line was established over a period of 2 months where MCF10A cells were subjected to increasing doses of AZD6738. Initially, naive parental cells were cultured in 1μM AZD6738 for 1 week until these cells underwent 4 population doublings. Then the inhibitor dose was increased to 2μM and again cells were left for 1 week until they underwent 4 population doublings. This culturing process was repeated again at 3, 4, and then 5μM concentrations until the cells could maintain consistent proliferation while continuously exposed to AZD6738 5μM. To confirm resistance, we compared the proliferation rate of these resistant cells to inhibitor-naive MCF10A cells at 5μM AZD6738. Importantly, the naive cells did not proliferate and slowly died off over a period of 1 week, while the resistant line continued to proliferate. For all experiments with the resistant cells, the ATR-R cells were relieved from ATR inhibition and cultured in media without any inhibitors for 24 hours prior to experimentation.

### Small Molecule Inhibitor Treatment

Small molecule inhibitors were dissolved in DMSO and added to cell culture medium to inhibit ATR, CHK1, CDK1, and CDK2. The following concentrations were used for inhibition: AZD6738 (ATRi, 5μM), VE822 (ATRi, 5μM), LY2603618 (CHK1i, 2μM), ChIR124 (CHK1i, 250nM), RO-3306 (CDK1i, 5μM), NU6140 (CDK2i, 1μM), KU60019 (ATMi, 5 μM), and NU7441 (DNA-PK, 5 μM). DMSO was used as a mock treatment for control groups. Hydroxyurea (RNRi, 1nM) was dissolved in H_2_O and thus H_2_O was used as a mock treatment for control groups.

Small Molecule Inhibitors

Small Molecule Inhibitors:
| **Name** | **Target** | **Source** | **Cat. Number** |
| --- | --- | --- | --- |
| AZD6738 | ATR | Selleck Chemicals | S7693 |
| VE822 | ATR | Selleck Chemicals | S7102 |
| LY2603618 | CHK1 | Selleck Chemicals | S2626 |
| ChIR124 | CHK1 | Selleck Chemicals | S2683 |
| RO-3306 | CDK1 | Selleck Chemicals | S7747 |
| NU6140 | CDK2 | Fisher Scientific | 33-011-0 |
| KU60019 | ATM | Fisher Scientific | S1570 |
| NU7441 | DNA-PK | Fisher Scientific | S2638 |
| Hydroxyurea | Ribonucleoside Reductase (RNR) | Sigma-Aldrich | H8627-1G |

### siRNA Transfection

Cells were seeded at 20,000 cells per well in glass-bottom 96-well plates (Cellvis P96-1.5P) and reverse-transfected with either non-targeting siRNA (siCTRL; Horizon Discovery, D-001810-10-05) or siRNA targeting cyclin B1 at a concentration of 20nM using DharmaFECT 1 (Horizon Discovery T-2005-01) following manufacturer’s guidelines. 20hrs post-transfection, the media was replaced with fresh growth media. Cells were fixed and analyzed 48hrs post-transfection.

siRNAs

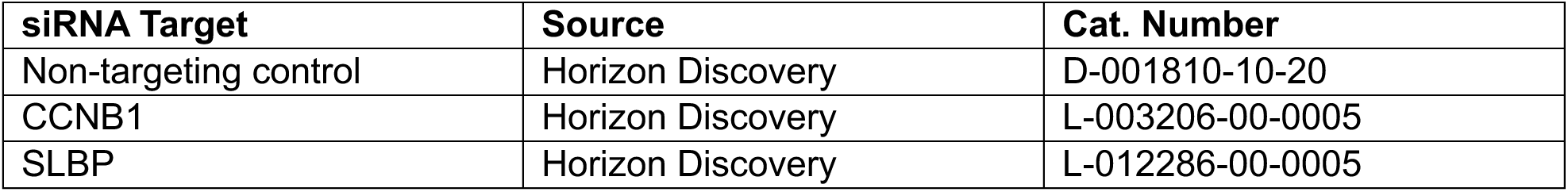

siRNAs:
| **siRNA Target** | **Source** | **Cat. Number** |
| --- | --- | --- |
| Non-targeting control | Horizon Discovery | D-001810-10-20 |
| CCNB1 | Horizon Discovery | L-003206-00-0005 |
| SLBP | Horizon Discovery | L-012286-00-0005 |

### Immunostaining

Cells were grown in 96-well plates with a glass-like polymer well-bottom (Cellvis, P96-1.5P). For cell cycle analysis, cells were treated with 10mM 5-ethynyl-2’-deoxyuridine (EdU) (Sigma-Aldrich. 900584) for 15 min prior to fixation unless otherwise stated. Pre-extraction was performed by incubating cells with 0.5% Triton X-100 (Millipore Sigma, T8787-50ML) in phosphate-buffered saline (PBS) for 1min and washing one time with 1x PBS before fixation. Cells were fixed with 4% paraformaldehyde (PFA) PBS for 10min, permeabilized with ice-cold methanol for 10min (even if pre-extracted), washed 1 time with 1X PBS, and blocked with 1% bovine serum albumin (BSA) in PBS for 30min at room temperature. Following blocking, cells were incubated with primary antibodies. All the primary antibodies were diluted in 1% BSA/PBS and incubated overnight with constant agitation at 4°C. After incubation with primary antibodies, cells were washed 3 times in 1X PBS and co-stained with DAPI (5μg/mL) and secondary antibodies. The secondary antibodies, anti-rabbit Alexa Fluor 488 conjugated antibody (Thermo Fisher Scientific, A-11008) and anti-mouse Alexa Fluor 568 conjugated antibody (Thermo Fisher Scientific, A-11004), were diluted 1:1000 in 1% BSA and incubated for 1hr at room temperature on a bench rocker. For EdU staining, the Click-iT reaction was performed following permeabilization using the Click-iT Cell Reaction Buffer kit (Invitrogen, C10269) and Alexa Fluor™ 647 Azide (Invitrogen, A10277) according to the manufacturer’s guidelines. Specifically, cells were washed with 3% BSA/PBS and incubated with the Click-iT reaction mixture with 1 µg/ml of Alexa Fluor™ 647 Azide for 30 min with at room temperature on a bench rocker. After the Click-iT reaction, cells were washed 2 times with 3% BSA/PBS, followed by one wash in 1X PBS. Finally, the 1x PBS was replaced once more before imaging.

Primary Antibodies

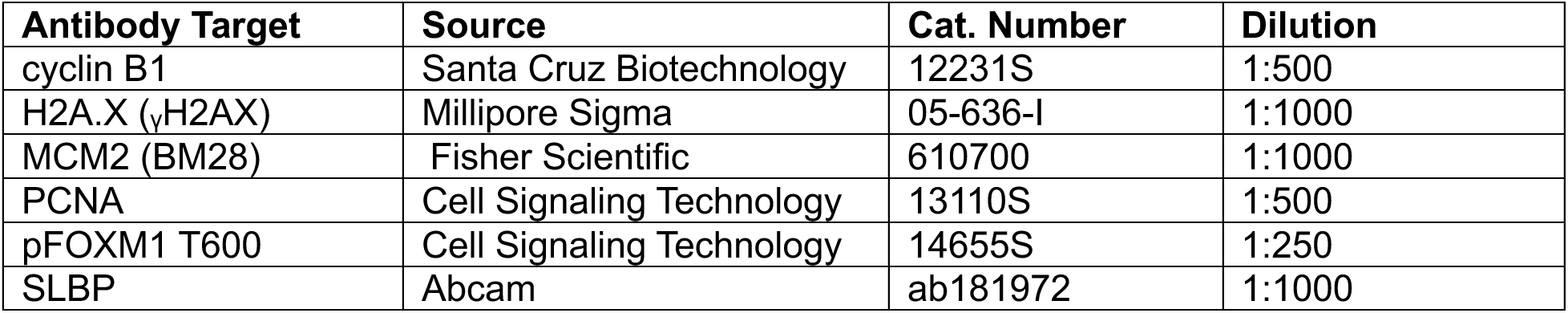

Primary Antibodies:
| Antibody Target | Source | Cat. Number | Dilution |
| --- | --- | --- | --- |
| cyclin B1 | Santa Cruz Biotechnology | 12231S | 1:500 |
| H2A.X (γH2AX) | Millipore Sigma | 05-636-I | 1:1000 |
| MCM2 (BM28) | Fisher Scientific | 610700 | 1:1000 |
| PCNA | Cell Signaling Technology | 13110S | 1:500 |
| pFOXM1 T600 | Cell Signaling Technology | 14655S | 1:250 |
| SLBP | Abcam | ab181972 | 1:1000 |

### Microscopy and Quantitative Image-Based Cytometry

Cells were imaged on a fully-automated ImageXpress Micro Confocal Imaging System (Molecular Devices). Widefield images were captured using a 20X Ph1S Plan Fluor ELWD ADM 0.45 NA objective and a sCMOS camera. Image analysis was performed using CellProfiler (Broad Institute). Intensity measurements were within a nuclear mask generated from DAPI-stained images. Background levels were determined for each fluorescence marker by assessing histogram plots of the signal intensities in each pixel across several images. The average background value across these images was then subtracted from the mean intensity of the given fluorescent marker in each cell. DNA content was determined using DAPI integrated intensity. Cytoplasmic cyclin B1 levels were determined by measuring the mean intensity within a ringed mask outside but adjacent to the nucleus. All other intensity measurements were within a nuclear mask generated from DAPI staining. Identification and quantification of mitotic slippage were derived from the analysis of DAPI-stained images. Color-coded scatter plots were generated using Tibco Spotfire.

## Data Availability

Data and images are available upon request.

## Acknowledgements

We thank members of the Saldivar laboratory for helpful discussions on the work presented in this paper. M.J.M is funded by an award from the Cancer Early Detection Advanced Research Center (CEDAR). J.C.S. is funded by NIH grant R35GM147710. The research reported in this publication used computational infrastructure supported by the Office of Research Infrastructure Programs, Office of the Director, of the National Institutes of Health under Award Number S10OD034224. The content is solely the responsibility of the authors and does not necessarily represent the official views of the National Institutes of Health.

## Author contributions

M.J.M. and J.C.S. conceived the study and wrote the paper. M.J.M. performed all experiments and analyses of the data. J.C.S. developed the ATRi-R cell line.

## Supplemental Materials

**Supplementary Fig. 1:**
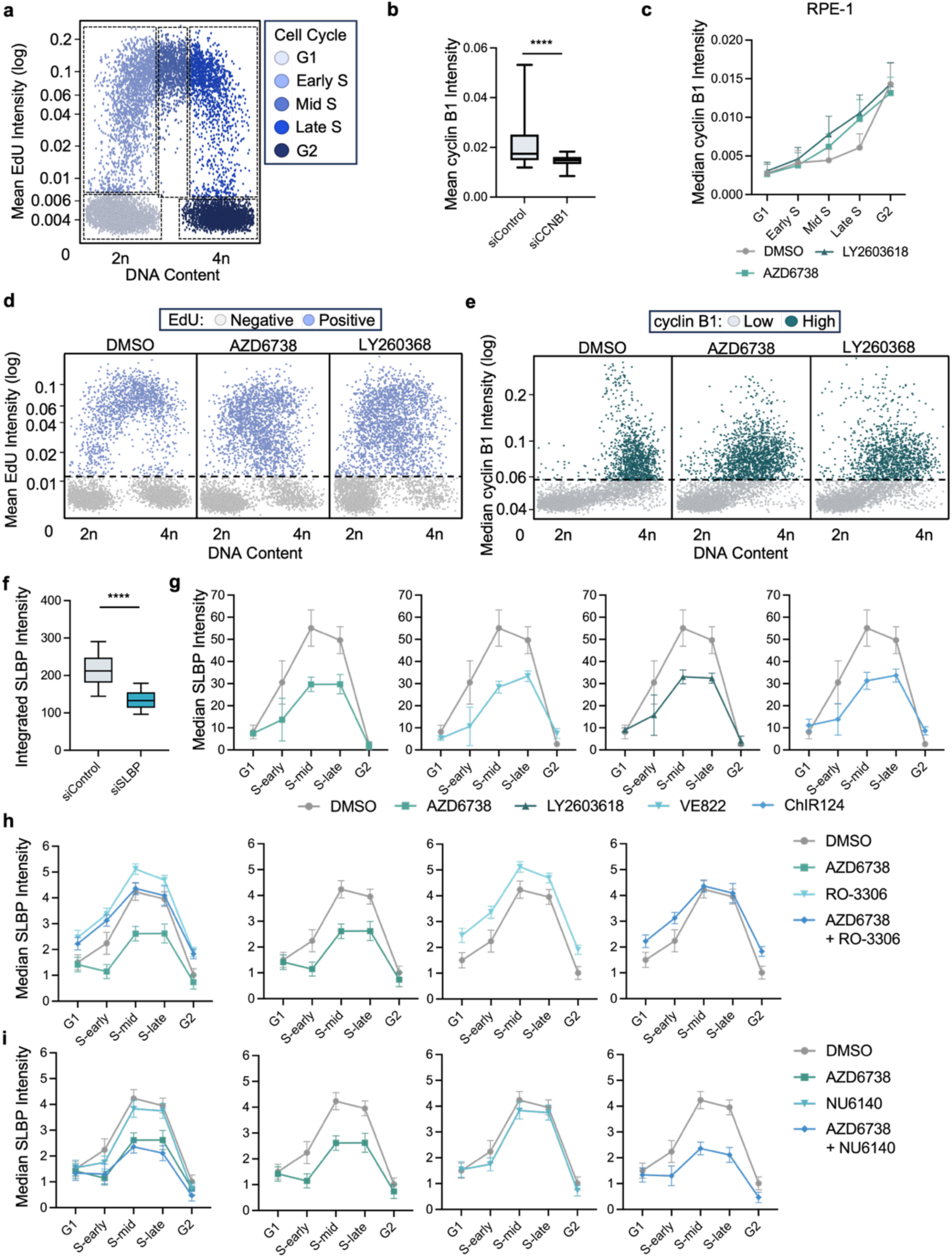
ATR regulates the S/G2 checkpoint in a CDK1-dependent manner. a.) Scatterplot of DNA content versus mean EdU intensity in MCF10A cells. Colors shown in the key and dotted lines denote gating used for analysis throughout this publication. b.) Scatterplot of DNA content versus median cyclin B1 intensity in MCF10A cells treated with DMSO, 5μM AZD6738, or 2μM LY2603618 for 16hrs. Colors shown in the key and dotted lines denote cyclin B1 gating used for analysis throughout this article. c.) Scatterplot of DNA content versus mean EdU intensity in MCF10A cells treated with DMSO, 5μM AZD6738, or 2μM LY2603618 for 16hrs. Colors shown in the key and dotted lines denote cyclin B1 gating used for analysis throughout this article. d.) Quantification of mean cyclin B1 intensity in MCF10A cells with non-targeting or cyclin B1 knockdowns to demonstrate cyclin B1 antibody specificity. A Mann-Whitney test was conducted to assess significance. Error bars are representative of 10-90 percentile range. P value: <0.0001. e.) Quantification of median cyclin B1 cytoplasmic intensity across the cell cycle from RPE-1 cells treated with DMSO, 5μM AZD6738, or 2μM LY2603618 for 16hrs (n=3 biological replicates). Data are presented as mean ± SEM. f.) Quantification of integrated SLBP intensity in MCF10A cells with non-targeting or SLBP knockdowns to demonstrate SLBP antibody specificity. Error bars are representative of 10-90 percentile range. A Mann-Whitney test was conducted to assess significance. P value: <0.0001. g.) Breakdown of Fig.1 quantification of median SLBP nuclear intensity across the cell cycle from into individual graphs. Data are presented as mean ± SEM. h.) Quantification of median SLBP nuclear intensity across the cell cycle from MCF10A cells treated with DMSO, 5μM AZD6738, 5μM RO-3306, or a combination of 5μM AZD6738 and 5μM RO-3306 for 1hr (n=3 biological replicates). Data are presented as mean ± SEM. i.) Quantification of median SLBP nuclear intensity across the cell cycle from MCF10A cells treated with DMSO, 5μM AZD6738, 1μM NU6140, or a combination of 5μM AZD6738 and 1μM NU6140 for 1hrs (n=3 biological replicates). Data are presented as mean ± SEM.

**Supplementary Fig. 2:**
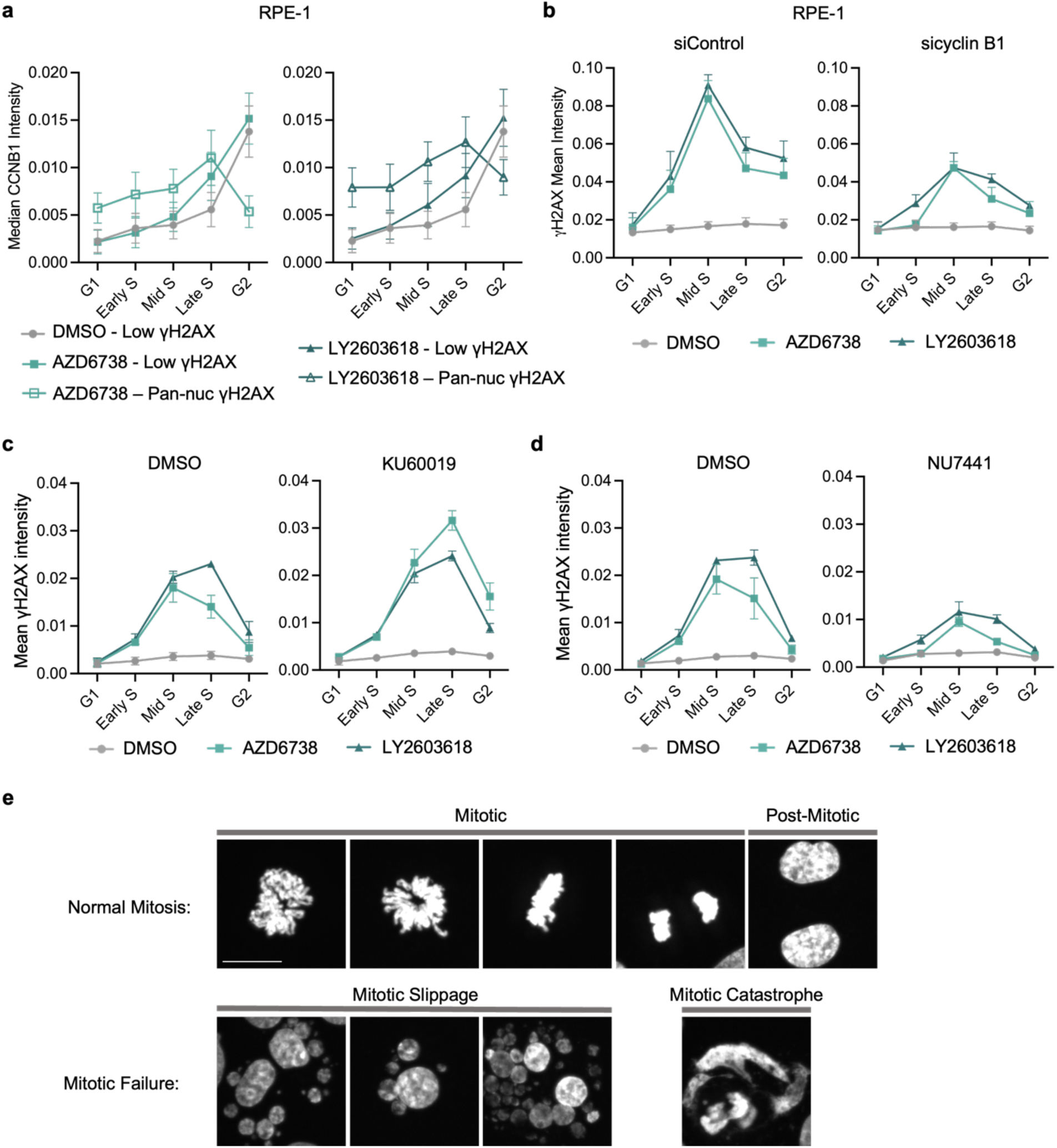
S/G2 checkpoint loss induces mitotic failure. a.) Quantification of median cyclin B1 cytoplasmic intensity in RPE-1 cells with low or high yH2AX across the cell cycle. Cells were treated with DMSO, 5μM AZD6738, or 2μM LY2603618 for 16hrs (n=3 biological replicates). Data are presented as mean ± SEM. b.) Quantification of mean yH2AX nuclear intensities across the cell cycle from control and cyclin B1 knockdown RPE-1 cells treated with DMSO, 5μM AZD6738, or 2μM LY2603618 for 16hrs (n=3 biological replicates). Data are presented as mean ± SEM. c.) Quantification of mean yH2AX nuclear intensity across the cell cycle from MCF10A cells treated with DMSO + DMSO, DMSO + 5μM AZD6738, DMSO + 2μM LY2603618, 5μM KU60019 + DMSO, 5μM KU60019 + 5μM AZD6738, 5μM KU60019 + or 2μM LY2603618 + 5μM KU60019 for 7hrs (n=3 biological replicates). Data are presented as mean ± SEM. d.) Quantification of mean yH2AX nuclear intensity across the cell cycle from MCF10A cells treated with DMSO + DMSO, DMSO + 5μM AZD6738, DMSO + 2μM LY2603618, 5μM KU60019 + DMSO, 5μM NU7441 + 5μM AZD6738, 5μM NU7441 + or 2μM LY2603618 + 5μM NU7441 for 7hrs (n=3 biological replicates). Data are presented as mean ± SEM. e.) Representative images of cells undergoing normal mitosis versus mitotic failure. Scale bar is equal to 15μm.

**Supplementary Fig. 3:**
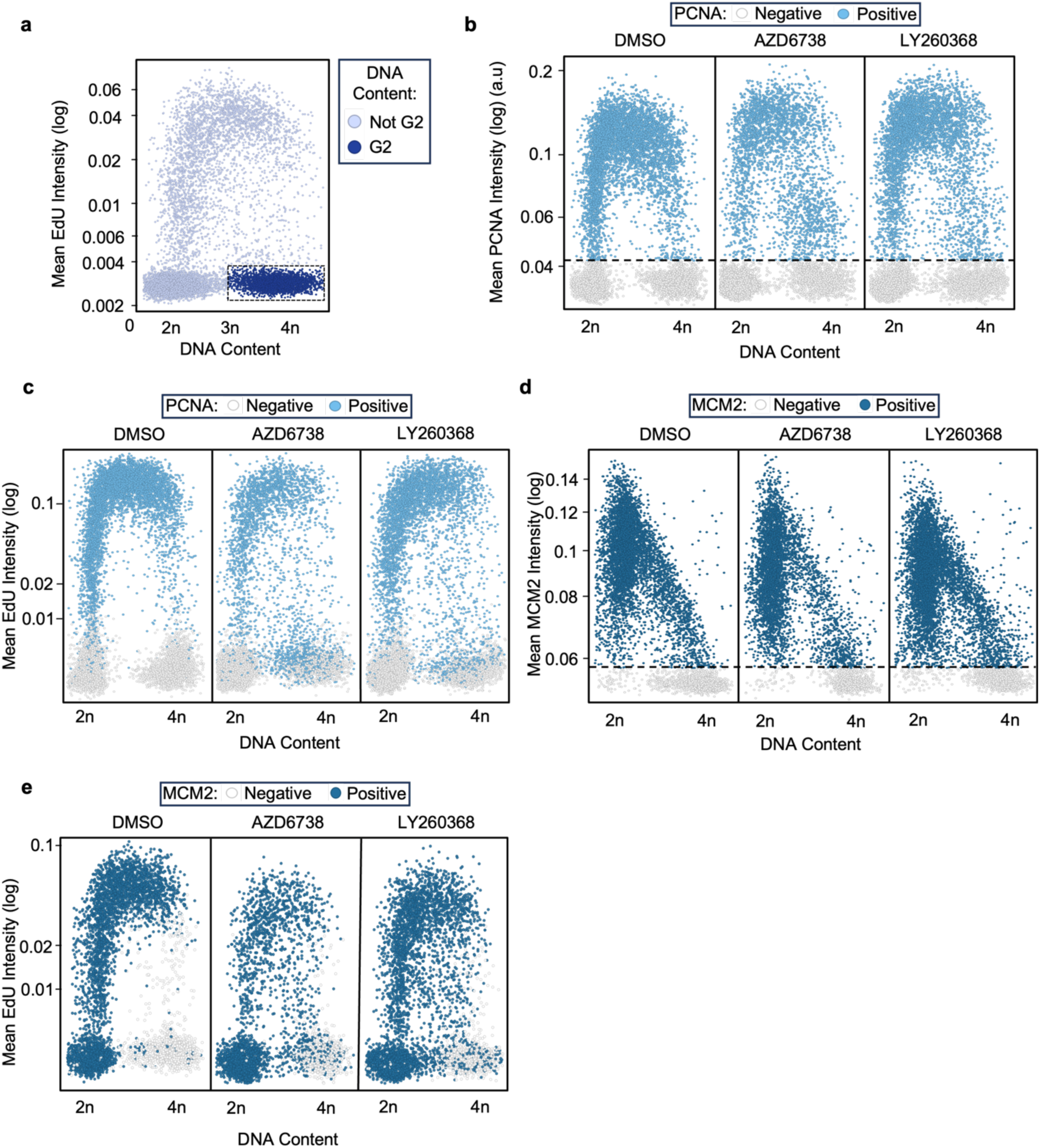
Gating thresholds for PCNA, MCM2, and EdU positive and negative cells. a.) Scatterplot of DNA content versus mean EdU intensity in MCF10A cells. Colors shown in the key and dotted lines denote gating of EdU negative cells with >3n DNA content (referred to as G2) used for analysis of chromatin-bound replication component retention. b.) Scatterplots of mean chromatin-bound PCNA intensity versus DNA content in pre-extracted MCF10A cells treated with DMSO, 5μM AZD6738, or 2μM LY2603618 for 16hrs. The dotted line and color of the points indicate the gating for PCNA positive and negative cells. c.) Scatterplots of mean EdU intensity versus DNA content in pre-extracted MCF10A cells treated with DMSO, 5μM AZD6738, or 2μM LY2603618 for 16hrs. The points are colored by status of being PCNA negative/positive as denoted by the associated key. Representative of n=3 biological replicates. d.) Scatterplots of mean chromatin-bound MCM2 intensity versus DNA content in pre-extracted MCF10A cells treated with DMSO, 5μM AZD6738, or 2μM LY2603618 for 16hrs. The dotted line and color of the points indicate the gating for MCM2 positive and negative cells. E.) Scatterplots of mean EdU intensity versus DNA content in pre-extracted MCF10A cells treated with DMSO, 5μM AZD6738, or 2μM LY2603618 for 16hrs. The points are colored by status of being MCM2 negative/positive as denoted by the associated key. Representative of n=3 biological replicates.

**Supplementary Fig. 4:**
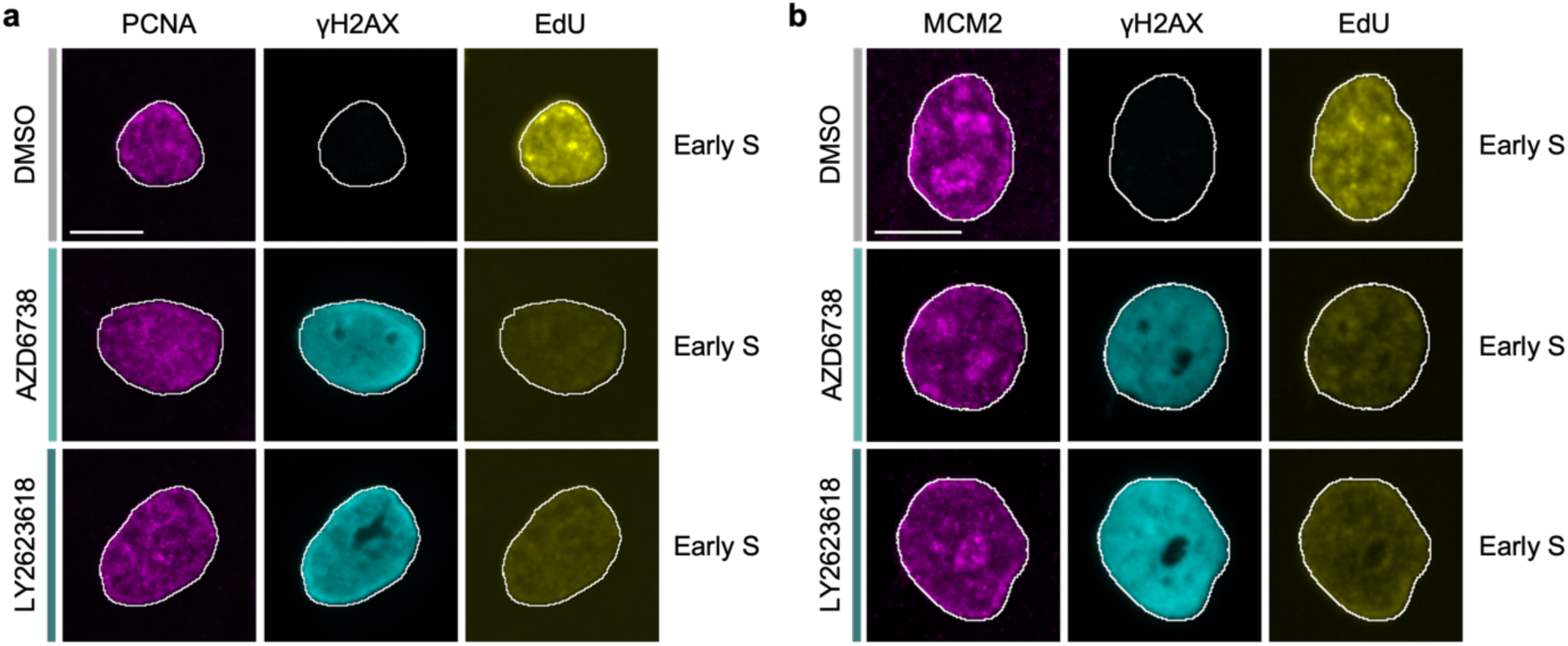
ATR pathway inhibition triggers replicative shutdown in early S phase. a.) Representative images of PCNA, yH2AX, and EdU in pre-extracted MCF10A nuclei treated with DMSO, 5μM AZD6738, or 2μM LY2603618 for 16hrs. Cell cycle status is indicated in the figure. Representative of n=3 biological replicates. Scale bar is equal to 10μm. b.) Representative images of MCM2, yH2AX, and EdU in pre-extracted MCF10A nuclei treated as in (a). Cell cycle status is indicated in the figure. Representative of n=3 biological replicates. Scale bar is equal to 10μm.

**Supplementary Fig. 5:**
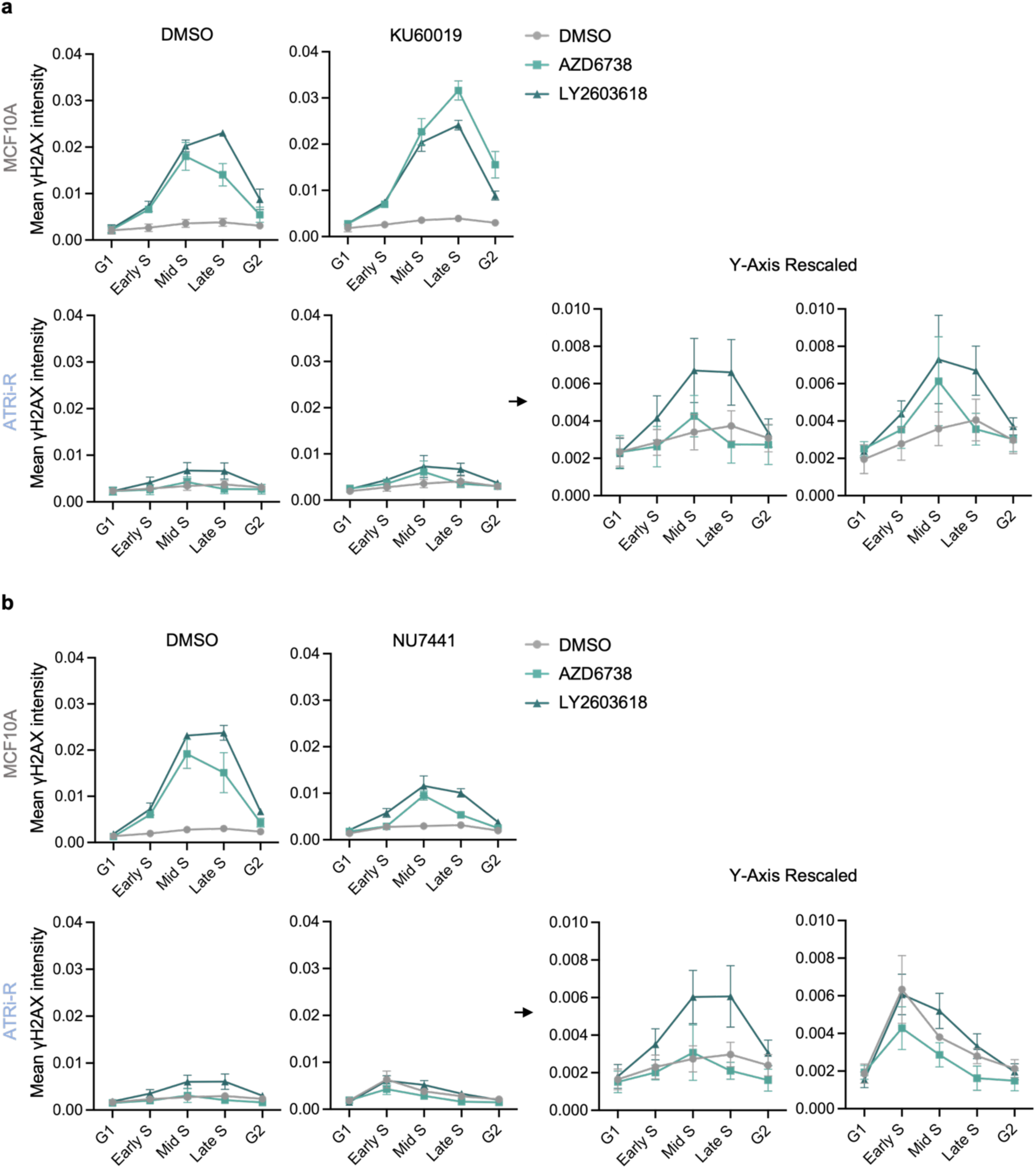
DNA-PK activity partially compensates for ATR in ATRi-R cells. a.) Quantification of mean yH2AX nuclear intensity across the cell cycle from ATRi-R cells treated with DMSO + DMSO, DMSO + 5μM AZD6738, DMSO + 2μM LY2603618, 5μM KU60019 + DMSO, 5μM KU60019 + 5μM AZD6738, 5μM KU60019 + or 2μM LY2603618 + 5μM KU60019 for 7hrs (n=3 biological replicates). Data from the parental MCF10A cell line is also presented in Supplementary Fig. 1c. Arrows indicate plots of ATRi-R values with scaled Y axis. Data are presented as mean ± SEM. d.) Quantification of mean yH2AX nuclear intensity across the cell cycle from ATRi-R cells treated with DMSO + DMSO, DMSO + 5μM AZD6738, DMSO + 2μM LY2603618, 5μM KU60019 + DMSO, 5μM NU7441 + 5μM AZD6738, 5μM NU7441 + or 2μM LY2603618 + 5μM NU7441 for 7hrs (n=3 biological replicates). Data from the parental MCF10A cell line is also presented in Supplementary Fig. 1d. Arrows indicate plots of ATRi-R values with scaled Y axis. Data are presented as mean ± SEM.

