## Supplemental Figures for "ATR enforcement of the S/G2 checkpoint prevents premature S phase shutdown and genome instability"

**Supplemental Materials**

### Figures

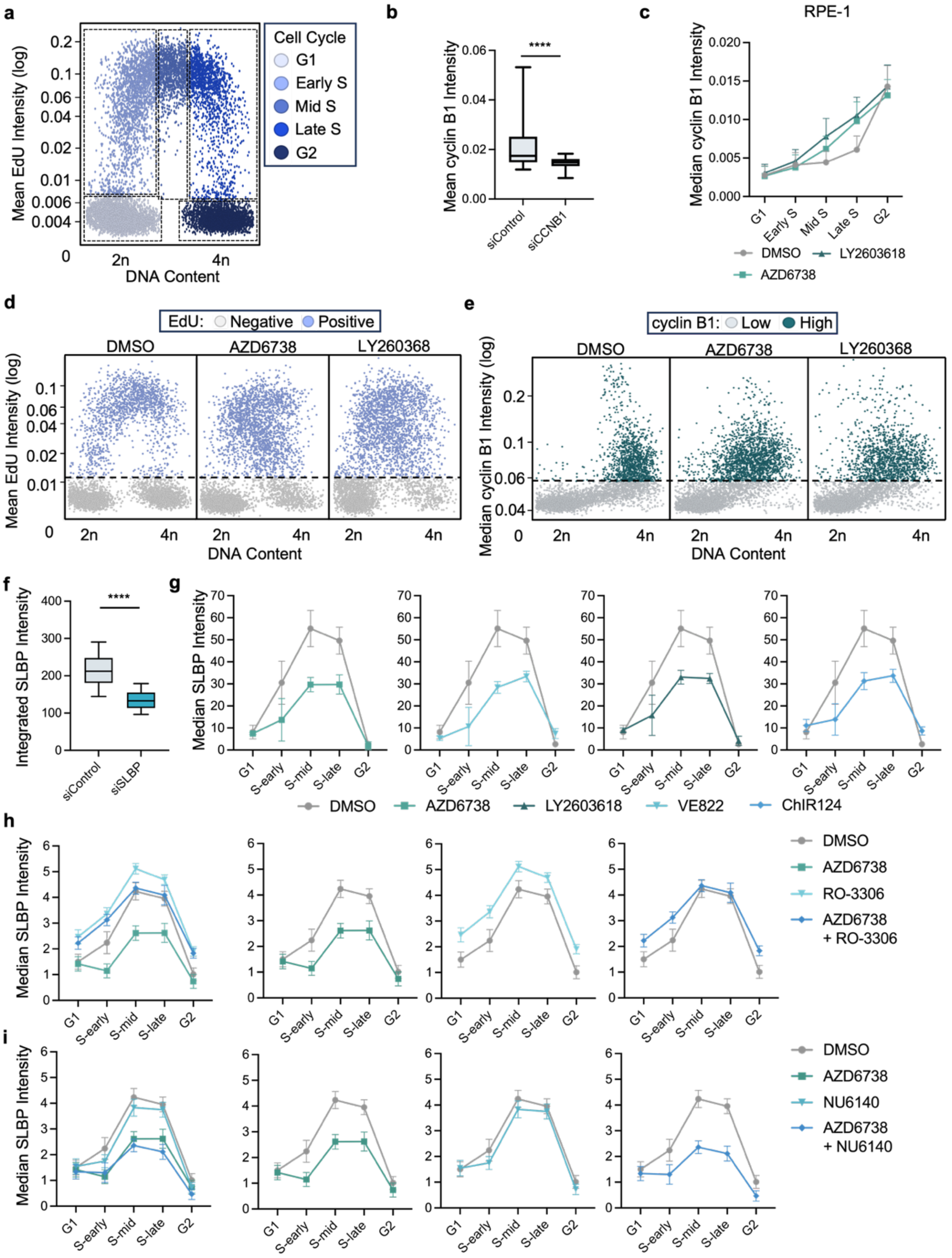


**Supplementary Fig. 1: ATR regulates the S/G2 checkpoint in a CDK1-dependent manner**

a.) Scatterplot of DNA content versus mean EdU intensity in MCF10A cells. Colors shown in the key and dotted lines denote gating used for analysis throughout this publication. b.) Scatterplot of DNA content versus median cyclin B1 intensity in MCF10A cells treated with DMSO, 5μM AZD6738, or 2μM LY2603618 for 16hrs. Colors shown in the key and dotted lines denote cyclin B1 gating used for analysis throughout this article. c.) Scatterplot of DNA content versus mean EdU intensity in MCF10A cells treated with DMSO, 5μM AZD6738, or 2μM LY2603618 for 16hrs. Colors shown in the key and dotted lines denote cyclin B1 gating used for analysis throughout this article. d.) Quantification of mean cyclin B1 intensity in MCF10A cells with non-targeting or cyclin B1 knockdowns to demonstrate cyclin B1 antibody specificity. A Mann-Whitney test was conducted to assess significance. Error bars are representative of 10-90 percentile range. P value: <0.0001. e.) Quantification of median cyclin B1 cytoplasmic intensity across the cell cycle from RPE-1 cells treated with DMSO, 5μM AZD6738, or 2μM LY2603618 for 16hrs (n=3 biological replicates). Data are presented as mean ± SEM. f.) Quantification of integrated SLBP intensity in MCF10A cells with non-targeting or SLBP knockdowns to demonstrate SLBP antibody specificity. Error bars are representative of 10-90 percentile range. A Mann-Whitney test was conducted to assess significance. P value: <0.0001. g.) Breakdown of Fig.1 quantification of median SLBP nuclear intensity across the cell cycle from into individual graphs. Data are presented as mean ± SEM. h.) Quantification of median SLBP nuclear intensity across the cell cycle from MCF10A cells treated with DMSO, 5μM AZD6738, 5μM RO-3306, or a combination of 5μM AZD6738 and 5μM RO-3306 for 1hr (n=3 biological replicates). Data are presented as mean ± SEM. i.) Quantification of median SLBP nuclear intensity across the cell cycle from MCF10A cells treated with DMSO, 5μM AZD6738, 1μM NU6140, or a combination of 5μM AZD6738 and 1μM NU6140 for 1hrs (n=3 biological replicates). Data are presented as mean ± SEM.

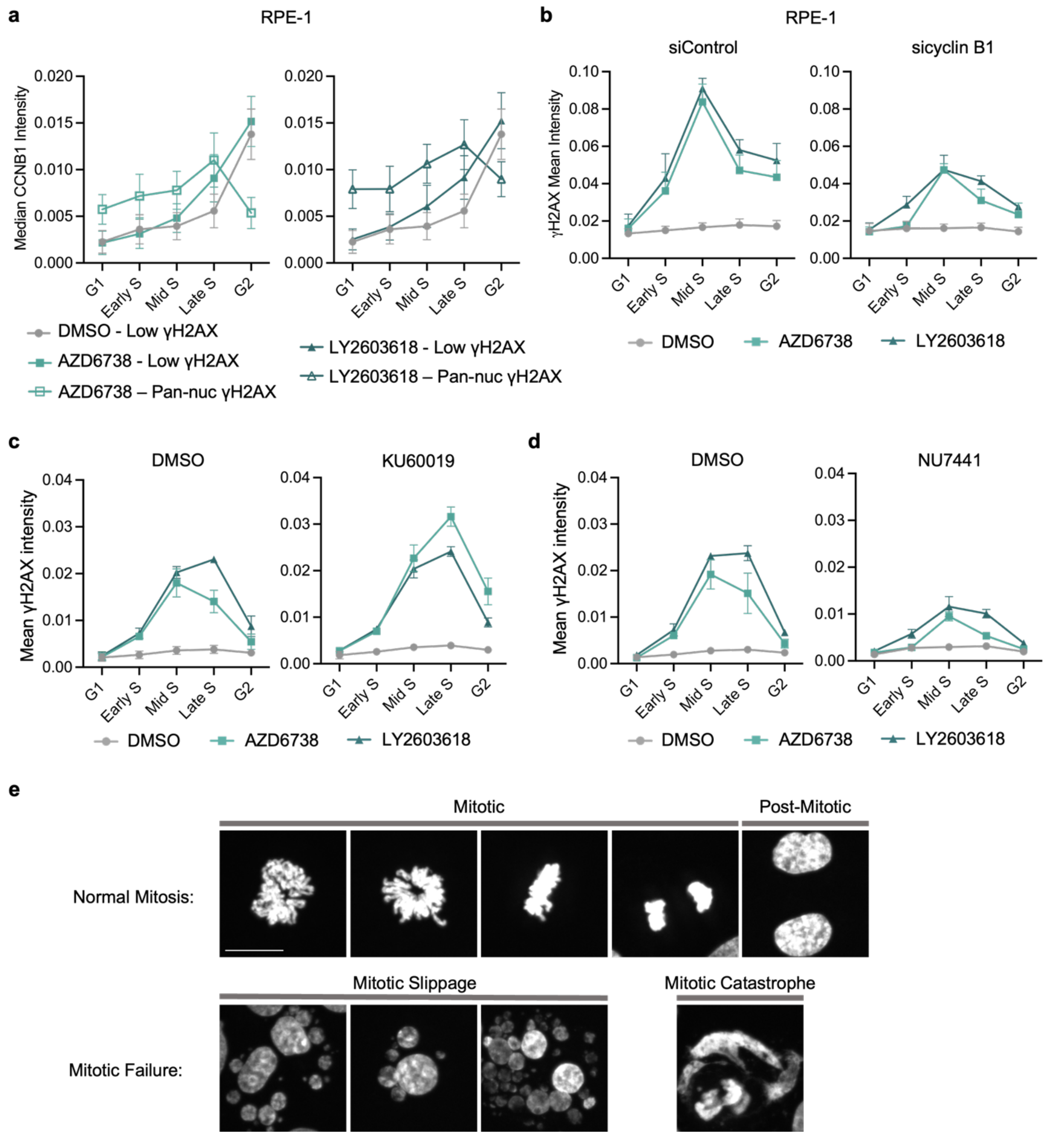


**Supplementary Fig. 2: S/G2 checkpoint loss induces mitotic failure**

a.) Quantification of median cyclin B1 cytoplasmic intensity in RPE-1 cells with low or high yH2AX across the cell cycle. Cells were treated with DMSO, 5μM AZD6738, or 2μM LY2603618 for 16hrs (n=3 biological replicates). Data are presented as mean ± SEM. b.) Quantification of mean yH2AX nuclear intensities across the cell cycle from control and cyclin B1 knockdown RPE-1 cells treated with DMSO, 5μM AZD6738, or 2μM LY2603618 for 16hrs (n=3 biological replicates). Data are presented as mean ± SEM. c.) Quantification of mean yH2AX nuclear intensity across the cell cycle from MCF10A cells treated with DMSO + DMSO, DMSO + 5μM AZD6738, DMSO + 2μM LY2603618, 5μM KU60019 + DMSO, 5μM KU60019 + 5μM AZD6738, 5μM KU60019 + or 2μM LY2603618 + 5μM KU60019 for 7hrs (n=3 biological replicates). Data are presented as mean ±

SEM. d.) Quantification of mean yH2AX nuclear intensity across the cell cycle from MCF10A cells treated with DMSO + DMSO, DMSO + 5μM AZD6738, DMSO + 2μM LY2603618, 5μM KU60019 + DMSO, 5μM NU7441 + 5μM AZD6738, 5μM NU7441 + or 2μM LY2603618 + 5μM NU7441 for 7hrs (n=3 biological replicates). Data are presented as mean ± SEM. e.) Representative images of cells undergoing normal mitosis versus mitotic failure. Scale bar is equal to 15μm.

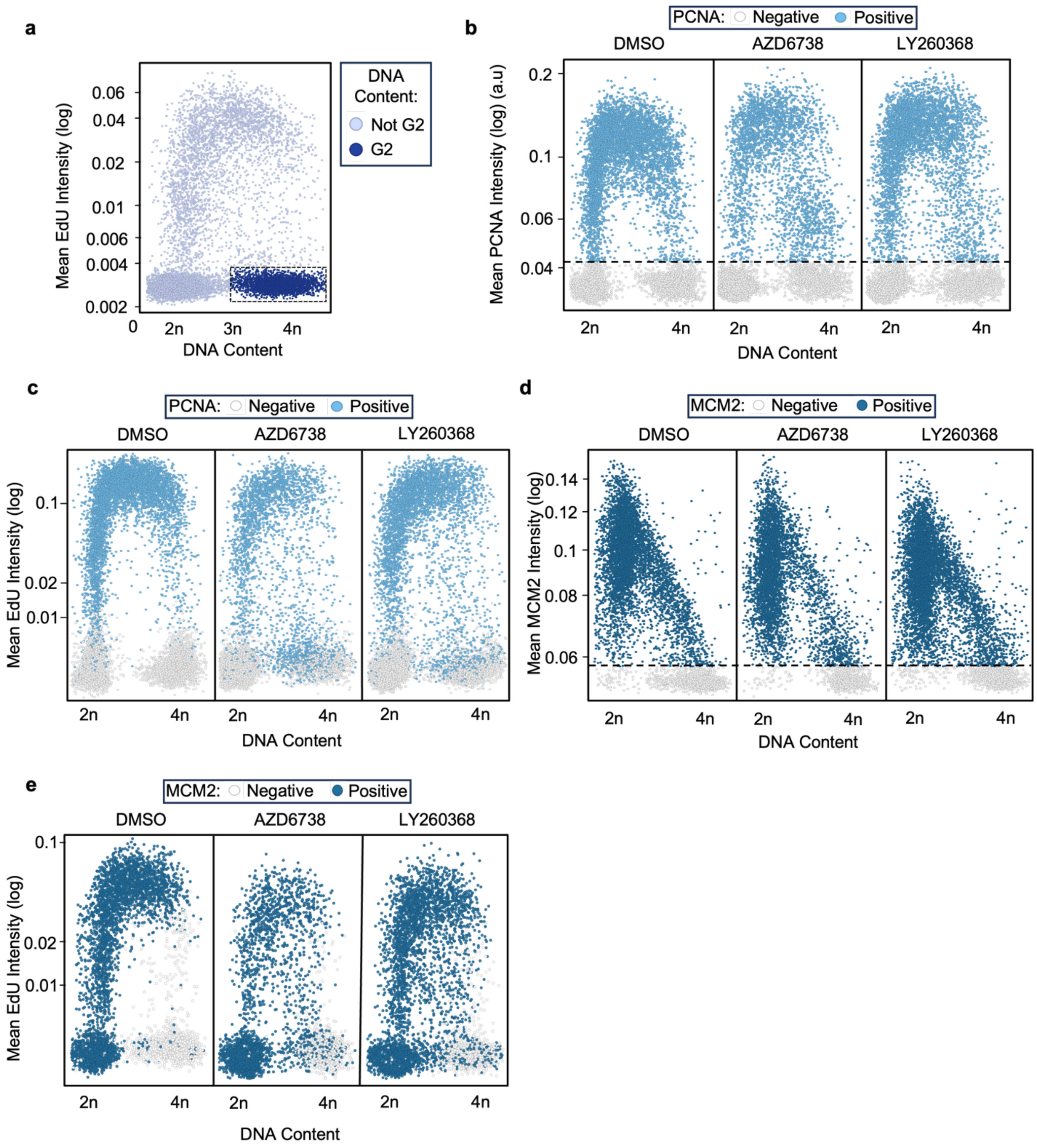


**Supplementary Fig. 3: Gating thresholds for PCNA, MCM2, and EdU positive and negative cells.**

a.) Scatterplot of DNA content versus mean EdU intensity in MCF10A cells. Colors shown in the key and dotted lines denote gating of EdU negative cells with >3n DNA content (referred to as G2) used for analysis of chromatin-bound replication component retention. b.) Scatterplots of mean chromatin-bound PCNA intensity versus DNA content in pre-extracted MCF10A cells treated with DMSO, 5µM AZD6738, or 2µM LY2603618 for 16hrs. The dotted line and color of the points indicate the gating for PCNA positive and negative cells. c.) Scatterplots of mean EdU intensity versus DNA content in pre-extracted MCF10A cells treated with DMSO, 5µM AZD6738, or 2μM LY2603618 for 16hrs. The points are colored by status of being PCNA negative/positive as denoted by the associated key. Representative of n=3 biological replicates. d.) Scatterplots of mean chromatin-bound MCM2 intensity versus DNA content in pre-extracted MCF10A cells treated with DMSO, 5μM AZD6738, or 2μM LY2603618 for 16hrs. The dotted line and color of the points indicate the gating for MCM2 positive and negative cells. E.) Scatterplots of mean EdU intensity versus DNA content in pre-extracted MCF10A cells treated with DMSO, 5μM AZD6738, or 2μM LY2603618 for 16hrs. The points are colored by status of being MCM2 negative/positive as denoted by the associated key. Representative of n=3 biological replicates.

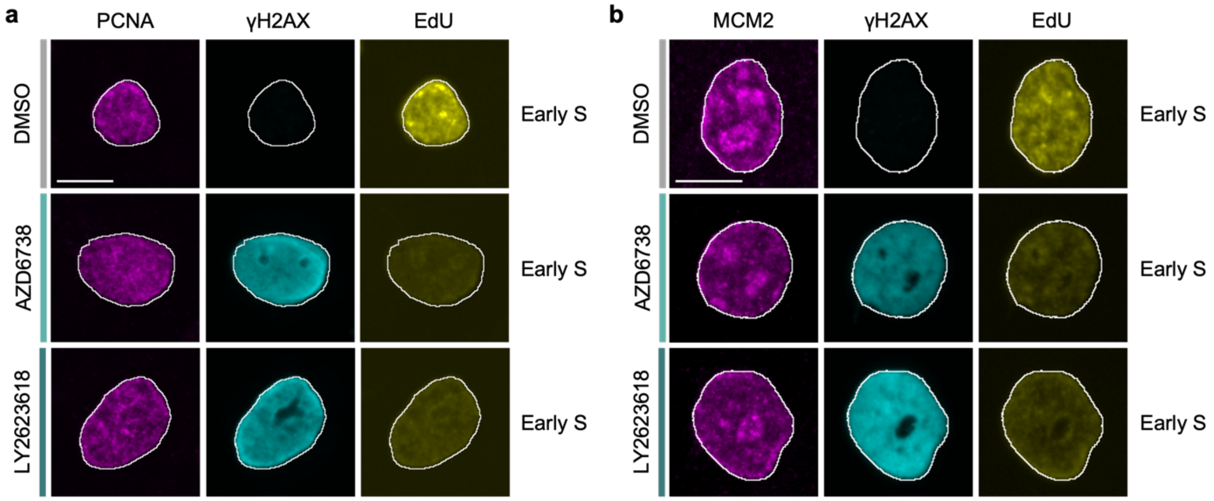


**Supplementary Fig. 4: ATR pathway inhibition triggers replicative shutdown in early S phase**

a.) Representative images of PCNA, yH2AX, and EdU in pre-extracted MCF10A nuclei treated with DMSO, 5µM AZD6738, or 2µM LY2603618 for 16hrs. Cell cycle status is indicated in the figure. Representative of n=3 biological replicates. Scale bar is equal to 10µm. b.) Representative images of MCM2, yH2AX, and EdU in pre-extracted MCF10A nuclei treated as in (a). Cell cycle status is indicated in the figure. Representative of n=3 biological replicates. Scale bar is equal to 10µm.

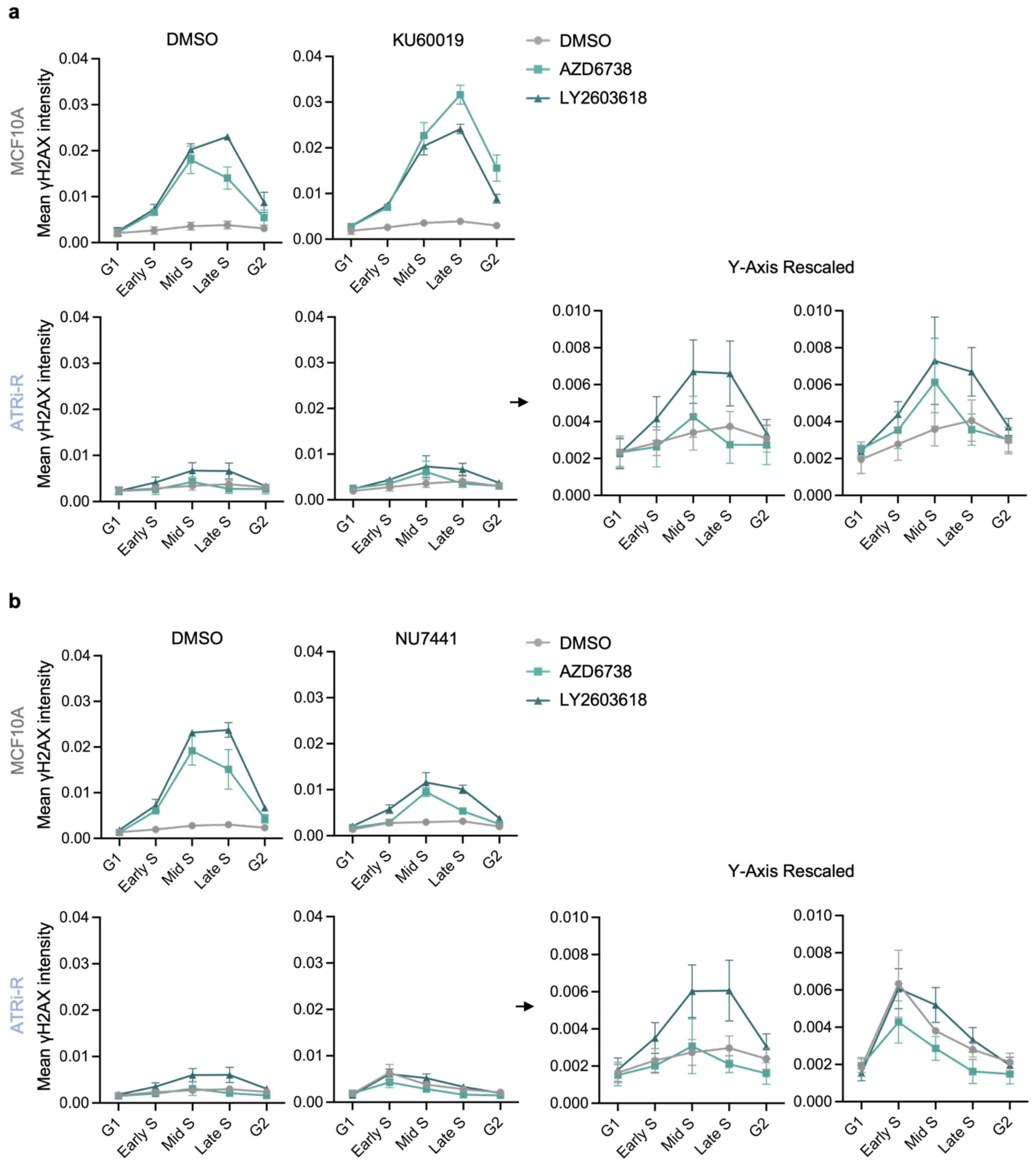


**Supplementary Fig. 5: DNA-PK activity partially compensates for ATR in ATRi-R cells**
a.) Quantification of mean γH2AX nuclear intensity across the cell cycle from ATRi-R cells treated with DMSO + DMSO, DMSO + 5μM AZD6738, DMSO + 2μM LY2603618, 5μM KU60019 + DMSO, 5μM KU60019 + 5μM AZD6738, 5μM KU60019 + or 2μM LY2603618 + 5μM KU60019 for 7hrs (n=3 biological replicates). Data from the parental MCF10A cell line is also presented in Supplementary Fig. 1c. Arrows indicate plots of ATRi-R values with scaled Y axis. Data are presented as mean ± SEM. d.) Quantification of mean γH2AX nuclear intensity across the cell cycle from ATRi-R cells treated with DMSO + DMSO, DMSO + 5μM AZD6738, DMSO + 2μM

LY2603618, 5µM KU60019 + DMSO, 5µM NU7441 + 5µM AZD6738, 5µM NU7441 + or 2µM LY2603618 + 5µM NU7441 for 7hrs (n=3 biological replicates). Data from the parental MCF10A cell line is also presented in Supplementary Fig. 1d. Arrows indicate plots of ATRi-R values with scaled Y axis. Data are presented as mean ± SEM.
